# A model for reticular dysgenesis shows impaired sensory organ development and hair cell regeneration linked to cellular stress

**DOI:** 10.1101/610204

**Authors:** Alberto Rissone, Erin Jimenez, Kevin Bishop, Blake Carrington, Claire Slevin, Stephen M. Wincovitch, Raman Sood, Fabio Candotti, Shawn M. Burgess

## Abstract

Mutations in the gene AK2 are responsible for Reticular Dysgenesis (RD), a rare and severe form of primary immunodeficiency in children. RD patients have a severely shortened life expectancy and without treatment die a few weeks after birth. The only available therapeutic option for RD is bone marrow transplantation. To gain insight into the pathophysiology of RD, we previously created zebrafish models for an AK2 deficiency. One of the clinical features of RD is hearing loss, but its pathology and causes have not been determined. In adult mammals, sensory hair cells of the inner ear do not regenerate; however, their regeneration has been observed in several non-mammalian vertebrates, including zebrafish. Therefore, we use our RD zebrafish models to determine if AK2 deficiency affects sensory organ development and/or hair cell regeneration. Our studies indicated that AK2 is required for the correct development, survival and regeneration of sensory hair cells. Interestingly, AK2 deficiency induces the expression of several oxidative stress markers and it triggers an increased level of cell death in the hair cells. Finally, we show that glutathione treatment can partially rescue hair cell development in the sensory organs in our RD models, pointing to the potential use of antioxidants as a supportive therapeutic modality for RD patients, not only to increase their chances of survival, but to prevent or ameliorate their sensorineural hearing deficits.

## Introduction

Reticular dysgenesis (RD) is a rare form of severe combined immunodeficiencies (SCID), a heterogenous group of immunological diseases usually characterized by profound defects to the T and B lymphoid lineages(*1*, *2*). In particular, RD patients also present a reduced number of neutrophils, with a specific block at the promyelocyte stage, and an unresponsiveness to GCSF stimulation in combination with severe sensorineural hearing loss(*3*, *4*). The immunodeficiencies associated with strong neutropenia leads these patients to recurrent severe infections and premature death within the first weeks or months of life. For these patients, a hematopoietic stem cell transplantation (HSCT) represents the only available treatment, but, because of the severity of the disease, pre-conditioning is necessary in order to increase the chances of engraftment of the donor stem cells(*5*). However, the HSCT does not improve the non-hematological defects such as the hearing loss, which has a huge impact on the cognitive functioning and quality of life of these children(*6*, *7*).

Mutations in the adenylate kinase 2 (AK2) gene have been shown to be responsible for the disease(*1*, *2*, *8*). AK2 is an enzyme localized to the intermembrane space of the mitochondria, where it plays an important role in sustaining mitochondrial respiration and cellular energy metabolism(*9*–*11*). Because the knockout of mouse AK2 showed an early embryonic lethality(*8*, *12*), other cellular and animal models needed to be developed. Insect models of AK2 deficiency indicated an essential role of the gene in embryonic growth and cell survival(*13*, *14*); moreover, they suggested that maternal AK2 mRNA can, at least initially, compensate for the lack of AK2 gene zygotic transcription. In zebrafish, AK2 knock-down induced by MO injection showed hematopoietic defects without affecting general embryonic development(*2*, *8*). These results were confirmed by two different mutant lines carrying a frameshift mutation in zebrafish AK2 exon 1 and a missense mutation in exon 4(*8*). Similar to what was observed *in vitro* in patient fibroblasts and CD34+ bone marrow cells(*2*, *11*), zebrafish mutants presented an increased level of cellular oxidative stress leading to apoptosis and cell death(*8*). Notably, these phenotypes can be reduced by the administration of antioxidants in both zebrafish mutants and, moreover, the same kind of treatment was able to rescue myeloid differentiation *in vitro* in induced pluripotent stem cells (iPSCs) obtained from fibroblasts of a RD patient(*8*).

Although most of the work linked the AK2 role to its bio-energetic activity, other evidence highlighted the presence of alternative functions, partially unrelated to its enzymatic activity and/or the mitochondrial localization(*5*). AK2 protein has been found to associate with dual-specificity phosphatase 26 (DUSP26), resulting in the suppression of cell proliferation by FADD dephosphorylation(*12*). In addition, it has been shown to be involved in an amplification loop that ensures the execution of the intrinsic apoptosis via an interaction with FADD and caspase 10(*15*). Previous reports showed that an AK2 deficiency impairs the regular induction of the unfolded protein response (UPR) mechanism *in vitro*(*16*, *17*). Finally, using RD patient-derived iPSCs, recent work showed a reduction of nuclear ATP levels in AK2-deficient cells during specific stages of hematopoietic differentiation(*10*). Reduced levels of nuclear ATP could be responsible for the altered transcriptional profile observed during hematopoietic differentiation(*10*, *11*). Overall all these studies collectively seem to suggest that, at least to some extent, the cellular AK2 roles can be cell-type or context specific.

Sensorineural hearing loss is the most common form of human hearing loss and it can be due to several different factors including genetic mutations, ototoxic compound exposure, aging, infectious diseases, or environmental stress, such as prolonged exposure to excessive noise(*6*, *18*). In general, all these different factors can induce damage to the mechanosensory hair cells in the organ of Corti or the stria vascularis and they can also impair the function of the spiral ganglion neurons or of the more proximal auditory structures(*19*). Because of the limited regenerative ability of mammals, hair cells cannot regenerate after damage and the resultant hearing loss is permanent. In contrast, non-mammalian vertebrates like zebrafish possess a huge regenerative potential and they can replenish hair cells during homeostasis or after damage, providing a model to study hair cells development and, in particular, hearing restoration after damage(*20*–*22*).

Although zebrafish inner ears lack a structure strictly equivalent to the mammalian cochlea, most of the functions present in all other vertebrates (i.e. hearing and vestibular functions) are fully conserved(*22*). Moreover, zebrafish possess another mechanosensory system on their body surface called the lateral line, which provides feedback information about water flow along the body of the fish(*23*–*25*). The lateral line is composed of a network of several small sense organs, the neuromasts, distributed over the zebrafish body in a specific pattern(*23*). It comprises two major branches, an anterior part that extends on the head (anterior lateral line or ALL), and a posterior part on the trunk and tail (PLL). Each neuromast consists of a central core of hair cells which are innervated by lateralis afferent neurons. The hair cells are then surrounded by two different types of non-sensory cells: supporting and mantle cells(*23*). Because of its structural simplicity and experimental accessibility, the lateral line, and in particular the PLL, has become a very popular model to study hair cells development and regeneration and it contributed significantly to the understanding of the molecular pathways that control those phenomena(*26*).

A previous study using adenylate kinase 2 immunostaining on mouse inner ears, indicated that AK2 protein localization was specifically limited to the lumen of the capillaries of the stria vascularis only in 7 days post-natal mouse embryos(*1*). Notably, no signal has been observed in other blood vessels or other cochlear structures such as the Organ of Corti(*1*). The stria vascularis is the portion of the inner ear responsible for the endolymph production, which is important for the metabolic support of the Organ of Corti(*1*, *19*). Therefore, it has been postulated that, in those specific structures, AK2 could work as an ecto-enzyme, controlling the ATP and/or potassium concentrations in the endolymph(*1*). As a consequence, its deficiency could result in hearing loss through a failure in the regulation of the potassium- and/ or ATP-homeostasis of the endolymph, as observed in the case of mouse connexin30 null mutants(*27*).

To shed light on the causes of RD hearing loss, we used the zebrafish models of RD(*8*). Our data showed that, zebrafish *ak2* is expressed in inner ear and neuromasts structures and its deficiency severely affects the survival of hair cells in those structures through an increased level of oxidative stress and cell death. Finally, the administration of glutathione (GSH) to zebrafish *ak2* mutants showed that antioxidant treatments were able to partially increase the number of hair cells in inner ear and PLL of *ak2*-deficient animals and this increase was both at the level of development and long-term survival of the hair cells.

## Results

### The Zebrafish *adenylate kinase 2* gene is expressed in sensory organs

Reticular dysgenesis is a rare hematological disorder caused by mutations in the adenylate kinase 2 (AK2) gene(*1*, *2*). Although at least one RD patient with skeletal defects has been reported so far(*28*), the only non-hematological clinical feature required for the diagnosis of RD is the sensorineural deafness/hearing disability(*4*, *5*). Because of the paucity of data, overall the pathophysiology of the hearing loss in RD patients is essentially uncharacterized. Here we used our zebrafish models of RD to better understand the physiopathology of hearing loss in RD patients. Initially we tested if the zebrafish AK2 gene (*ak2*) was expressed in sensory organs such as the inner ear or the lateral line system. Whole mount in situ hybridization (WISH) analysis of endogenous *ak2* expression during zebrafish development showed that from 3 days post fertilization (dpf) *ak2* gene is expressed in different anatomical regions, included the otic vesicle (Figure 1A). To better characterize the territories of *ak2* expression in the otic vesicle, we performed transversal sectioning on 5 dpf stained embryos. As shown in Figure 1-figure supplement 1, positive cells were observed in different structures of the inner ear such as the cristae and the anterior and posterior maculae. Although we were not able to observe the *ak2* expression in neuromasts in whole-mount staining, probably due to the low levels of expression, notably, in the same transverse sections we also found positive cells inside the neuromasts of the anterior lateral line (ALL) (Figure 1-figure supplement 1, black arrows). Based on their rounded morphology and large nuclei, in contrast to the slim and elongated cell bodies of the supporting cells(*29*), those cells represent putative hair cells.

**Figure 1.**
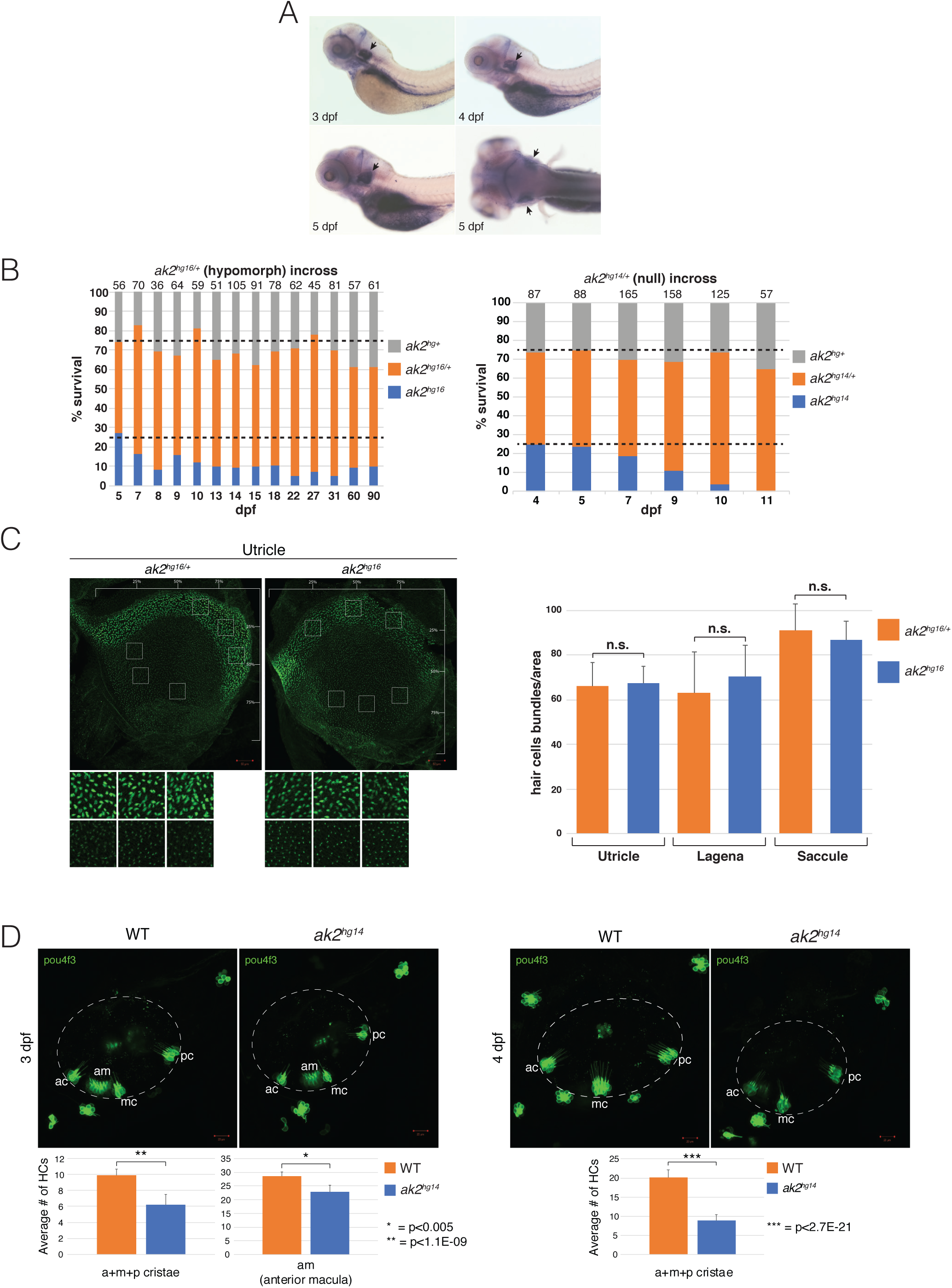
*ak2* larval expression in the otic vesicle and sensory phenotypes of the zebrafish *ak2* alleles. (A) Whole-mount in situ hybridization on wild-type zebrafish larvae at different developmental stages (3-5 dpf). Lateral (3-5 dpf) and dorsal (5 dpf) views with anterior to the left. Black arrows point to *ak2* expression in the otic vesicles. (B) Analysis of the survival rates for wild-type, *ak2* heterozygous, and *ak2* mutant larvae at different days post fertilization (dpf). Dashed lines indicate the expected ratio following Mendelian inheritance. (C) Comparison between hair cell bundle density in utricles of *ak2^hg16/+^* and *ak2^hg16^* adult fish. Left panel: the hair cell bundles were stained with phalloidin (green). In each structure, hair cell bundle counts were sampled within 2500 µm^2^ areas indicated by white boxes and insets below each figure. Scale bars: 50 µm. Right panel: quantification of hair cell bundle density in *ak2^16/+^* and *ak2^16^* animals. (D) Comparison of confocal maximum projections of inner ear regions (dashed circles) from Tg(*pou4f3*:GAP-GFP) wild-type (WT) and *ak2^hg14^* mutants at 3 and 4 dpf. Scale bars: 20 µm. Graphs below each set of pictures show the comparison of the average number of GFP positive hair cells in the anterior macula (3 dpf) and cristae (3 and 4 dpf) between *ak2*-deficient animals and controls (WT). Asterisks indicate statistically significant differences. c: crista; pm: posterior macula; n: neuromast; ac: anterior crista; mc: medial crista; pc: posterior crista; am: anterior macula. The error bars indicate standard errors; p indicates the p-values, according to Student’s t-test. N.s.: not significant according to Student’s t-test.

These data suggest that, in zebrafish, *ak2* is expressed in sensory organs and that the lack of its expression could potentially affect inner ear or neuromast development to some extent.

### Lack of *ak2* affects inner ear hair cell development

In a previous study, we showed that two different *ak2* mutants, carrying a missense mutation (previously denoted as *ak2^L124P/L124P^*, and here indicated as *ak2^hg16^*) or frame-shift mutations in exon1 of *ak2* (*ak2^del2/del2^* and *ak2^ins4/ins4^* mutants, here designated as *ak2^hg14^* and *ak2^hg15^*, respectively), presented very similar severe hematopoietic phenotypes affecting hematopoietic stem and precursor cell development(*8*). However, survival analysis showed that some *ak2^hg16^* mutants can reach adulthood, although with a lower than expected ratio of survivors (~10% compared to the expected percentage of ~25%) (Figure 1B, left panel). In contrast, *ak2*-deficient mutants *ak2^hg14^* and *ak2^hg15^*, lacking the *ak2* mRNAs(*8*), start to die around 5 dpf; and by 11 dpf we were not able to find any surviving homozygous mutants (Figure 1B, right panel and data not shown). These data suggest that, despite the severity of the hematopoietic defects observed in all *ak2* mutants, *ak2^hg16^* likely represents a hypomorphic mutation, while the other two alleles (*ak2^hg14^* and *ak2^hg15^*) are more likely fully null. Notably, in *ak2^hg16^* adults we did not observe any behavioral signs indicating potential sensorineural defects(*22*, *30*) such as “circling” or the lack of a startle response. When we compared hair cell bundle density of inner ear structures from *ak2^hg16^* and their heterozygous siblings (*ak2^hg16/+^*) of the same age, we did not find a statistically significant difference (Figure 1C and Figure 1-figure supplement 2). However, using the Tg(*pou4f3*:GAP-GFP) transgenic line to visualize the mature hair cells, we found a statistically significant difference in the average number of hair cells in 3 and 4 dpf developing inner ears of *ak2^hg14^* mutants compared to their control siblings (Figure 1D).

Taken together, these data suggest that a more severe *ak2* deficiency is needed to impair the correct development of the hair cells in the zebrafish inner ears compared to the hematopoietic defects seen in the hypomorphic allele.

### *ak2* deficiency affects posterior lateral line development

One hypothesis for the sensorineural hearing loss in AK2-deficient patients is that the composition of the luminal space is compromised, resulting in hair cell loss(*1*, *5*). To understand if hair cells in a different context (i.e. not in a luminal space) could similarly be affected by the lack of *ak2* expression, we examined the posterior lateral line (PLL) of our knockout fish. In zebrafish larvae, the PLL possess at least three different populations of neuromasts denoted as primary, secondary and intercalary (IC)(*24*). Primary neuromasts are deposited by the first migrating primordium (primI) which travels from the otic vesicle along the midline of the trunk on both sides of the embryo(*23*, *31*, *32*) to the tail, regularly depositing primordial neuromasts. Moreover, all the primary neuromasts are connected by a thin continuous stripe of cells (interneuromast cells or INCs) which, later during development, act as multipotent stem cells to produce intercalary neuromasts(*33*). By 2 dpf, the migration of the first primordium is complete as it reaches the most caudal part of the trunk of the embryos. At approximately the same time, a second, slower primordium (primII) starts its migration from the otic vesicle following a similar route used by the primI and it generates a second set of neuromasts called “secondary neuromasts” (or LII)(*24*). In *ak2^hg16^* hypomorphic embryos we did not observe any major defects to the PLL formation as indicated by alkaline phosphatase (AP) staining(*34*) at 4-5 dpf or Yo-PRO-1 staining(*35*, *36*) of functional hair cells from 3 to 5 dpf (Figure 2-figure supplement 1A and 1B, respectively). Additionally, we did not observe any statistical difference in the regenerative ability of hair cells after copper sulfate ablation(*37*, *38*) (Figure 2-figure supplement 1C). In contrast, from ~3 dpf *ak2^hg14^* embryos presented a lack of positive AP signal and Yo-PRO-1 staining in secondary neuromasts (LII.1-3) (Figure 2A and 2B, respectively), suggesting late defects to secondary neuromast formation. Notably, although the overall development of primary neuromasts appeared normal in *ak2^hg14^* embryos (Figure 2A), further analysis showed a reduction in the average number of Yo-PRO-1^+^ hair cells/neuromast after 3 dpf (Figure 2C), indicating measurable defects during primary neuromast development and maturation. To determine if loss of *ak2* function has a negative impact on the regeneration of hair cells, we checked if *ak2* expression was present after hair cells ablation with copper sulfate. WISH analysis performed at different time points after complete hair cell ablation with a 1-hour treatment with 10 µM copper sulfate, showed that *ak2* expression was increased in neuromasts 3 hours post ablation (hpa) (data not shown). Accordingly, we also observed a strong reduction of Yo-PRO-1^+^ hair cells in *ak2^hg14^* embryos at 2 days post ablation (dpa) compared to control siblings (Figure 2D). Moreover, we observed similar results performing copper sulfate hair cell ablation assay on *ak2^hg14^* in Tg(*pou4f3*:GAP-GFP) or Tg(*tnks1bp1*:EGFP;*atoh1a*:dTOM) backgrounds which label hair cells and supporting cells of the neuromasts (Figure 2-figure supplement 2A and 2B, respectively) with a strong reduction of the number of both mature *(pou4f3*:GAP-GFP^+^) and immature (*atoh1a*:dTOM^+^) hair cells in primary neuromasts.

**Figure 2.**
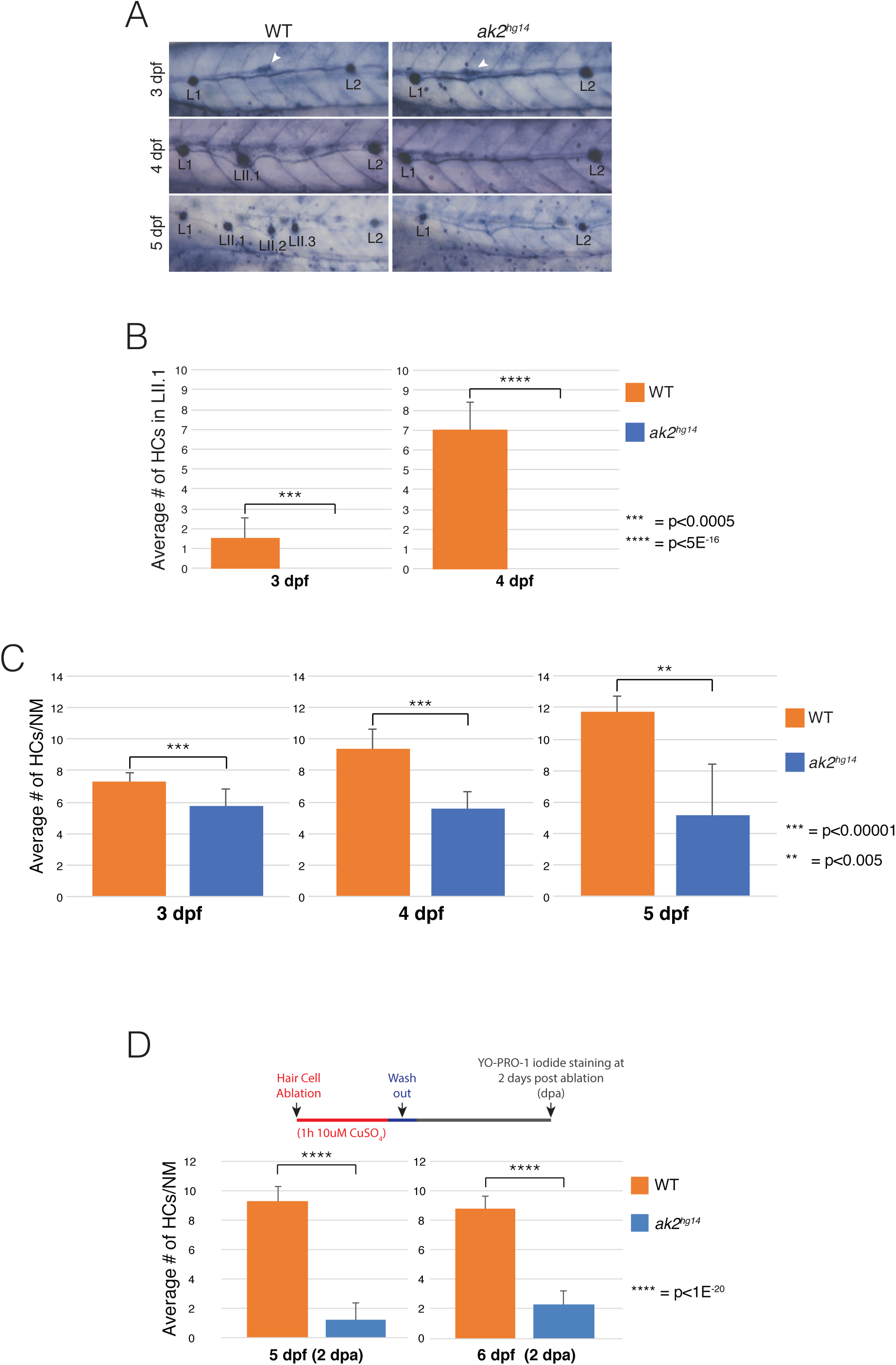
Reduced hair cell number and secondary neuromast differentiation phenotypes in *ak2^hg14^* mutants. (A) Lateral views of the trunk of 3-5 dpf *ak2^hg14^* embryos and their siblings stained for endogenous alkaline phosphatase activity. White arrowheads indicate the migrating secondary primordium (primII). LII neuromasts are severely reduced in number in *ak2* mutants. (B) Yo-PRO-1 iodide staining of HCs in lateral line Neuromasts to compare the average number of HCs in LII.1 secondary neuromasts in *ak2^hg14^* embryos and their wild-type siblings during development. (C) Analysis of the average number of HCs per neuromast in Yo-PRO-1 iodide stained *ak2^hg14^* embryos and their siblings during development. (D) Analysis of the average number of HCs per neuromast in Yo-PRO-1 iodide stained *ak2^hg14^* embryos and their siblings after 2 days post ablation (dpa) with CuSO_4_. The error bars indicate standard errors. Asterisks denote significant values according to Student’s t-test. p indicates the p-values. L1, L2 and L3 designate primary Neuromasts; LII.1, LII.2 and LII.3 indicate the lateral line secondary neuromasts.

Overall these data suggest that lack of *ak2* severely impairs the correct development and maturation of the secondary neuromasts, partially affecting the development and the regenerative ability of the hair cells of the primary neuromasts as well, leading us to hypothesize that hair cell and neuromast precursor cells are being prematurely depleted in *ak2* deficient animals. The observed deficiencies in lateral line hair cells which are on the skin surface also suggests the hypothesis of luminal composition deficiencies in humans may not be correct.

### *ak2* deficiency affects correct expression of neuromast markers in secondary neuromasts

To better characterize the PLL phenotypes observed in *ak2*-deficient mutants, we perform WISH and confocal microscope analyses of *ak2*-deficient (*ak2^hg14^* and *ak2^hg15^*) embryos in wild-type and different transgenic backgrounds. WISH analysis with the immature hair cell marker *atoh1a* showed a specific reduction of expression in primary neuromasts (L1 and L2) and a complete lack of expression in secondary neuromasts of *ak2^hg14^* embryos at 5 dpf, but not in control siblings (Figure 3A and Figure 3-figure supplement 1A) or in the *ak2^hg16^* hypomorphic mutants (Figure 3B). Interestingly, WISH analysis with the *eya1* marker labeling primII and lateral line neuromasts(*29*) showed no noticeable defects until 3.5 dpf (Figure 3C and Figure 3-figure supplement 1B). However, from ~4 dpf *eya1* expression was undetectable in the secondary primordium (Figure 3C) and strongly reduced or completely absent in LII.1 and LII.2 neuromasts (Figure 3C).

**Figure 3.**
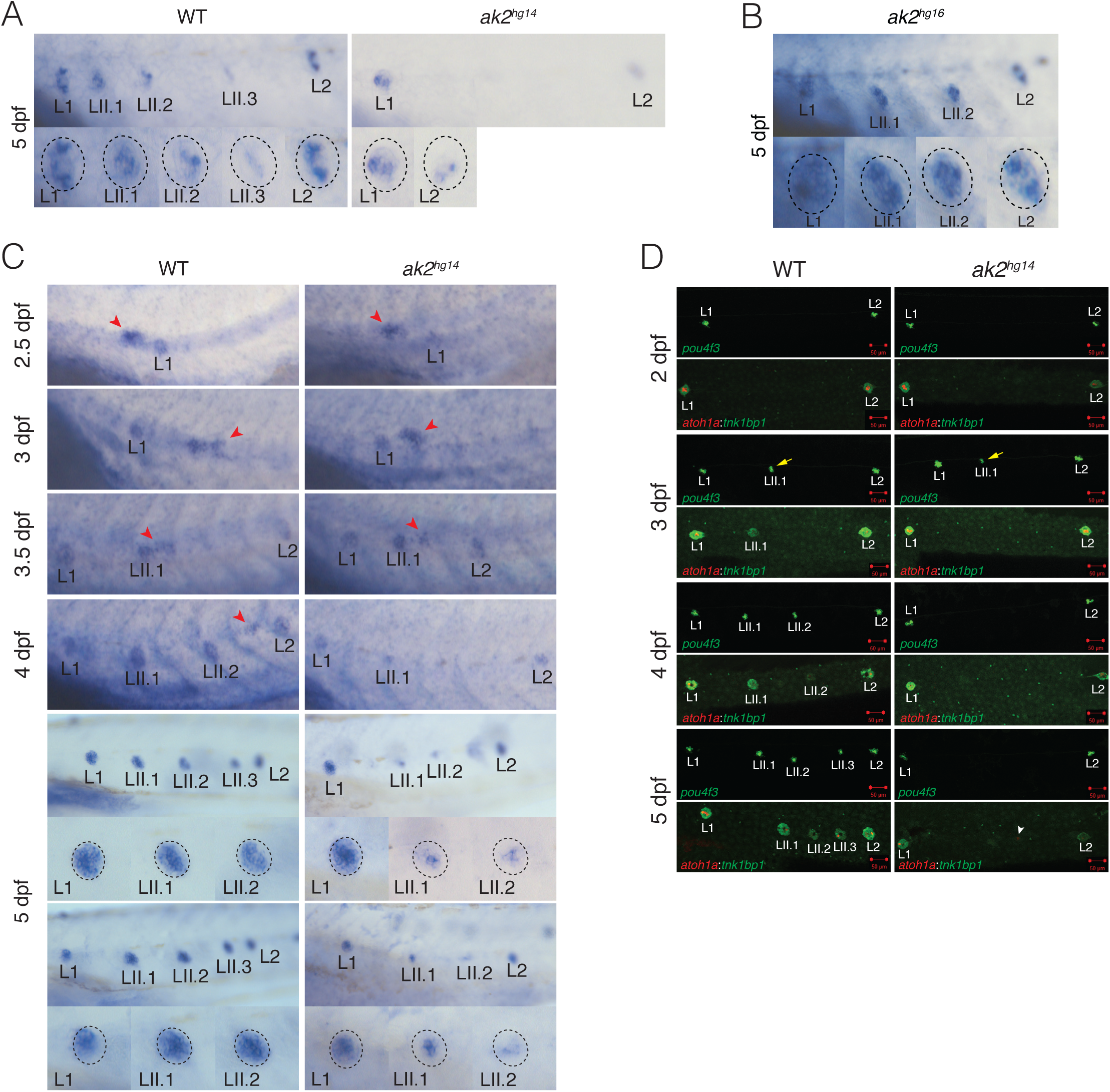
Characterization of PLL neuromast markers in lateral line of *ak2^hg14^* mutants. (A) WISH analysis of *atoh1a* marker expression in primary and secondary neuromasts on 5 dpf *ak2^hg14^* embryos and their siblings. (B) *atoh1a* expression analysis in primary and secondary neuromasts of 5 dpf *ak2^hg16^* embryos. (C) WISH analysis of *eya1* expression from 2.5 to 5 dpf in *ak2^hg14^* embryos and their siblings. At 2.5 and 3 dpf, a trunk region around L1 is shown; at 3.5 and 4 dpf pictures show the trunk region between L1 and L2 primary neuromasts. Red arrowheads indicate the migrating secondary primordium (primII). Each panel shows higher magnification of the regions around each primary and secondary neuromast from corresponding pictures in Figure 3 supplement 1. (D) Confocal analysis from 2 to 5 dpf of trunk regions of *ak2^hg14^* embryos and their siblings (WT) with different transgenic backgrounds to visualize components of the lateral line rosettes. *pou4f3* denotes the Tg(*pou4f3*:GAP-GFP) line labelling the mature HCs (green); *atoh1a*:*tnks1bp1* indicates the Tg(*tnks1bp1*:EGFP; *atoh1a*:dTOM) line labelling immature HCs (red) and supporting cells (green), respectively. Scale bars: 50 µm. Yellow arrows indicate *pou4f3*:GFP^+^ HCs in secondary neuromasts; white arrow points to solitary *atoh1a:*dTOM*^+^* cell. L1 and L2 define primary neuromasts; LII.1, LII.2 and LII.3 designate the lateral line secondary neuromasts.

For a more detailed study of the secondary neuromast formation and development in *ak2*-deficient embryos, we crossed *ak2^hg14/+^* and *ak2^hg15/+^* animals with transgenic lines marking different cellular components of neuromasts. As previously observed using Yo-PRO-1 staining (Figure 2B), the analysis of *ak2^hg14^* Tg(*pou4f3*:GAP-GFP) embryos further confirmed the lack of GFP^+^ cells (corresponding to hair cells) in secondary neuromasts from ~4 dpf (Figure 3D). Notably, at 3dpf the confocal analysis showed the presence of GFP^+^ cells in LII.1 neuromasts, although with a severely reduced number of cells compared to the controls (Figure 3D, yellow arrows). The lack of Yo-PRO-1 positive cells in LII.1 of *ak2* homozygous mutants at the same stage (Figure 2B) suggested these GFP positive hair cells are not fully mature and functional, despite expressing *pou4f3*.

Using the double transgenic line Tg(*tnks1bp1*:EGFP; *atoh1a*:dTOM)(*39*) to label immature hair cells (red) and supporting cells (green), we were able to highlight further defects in secondary neuromast formation (Figure 3D). In particular, the analysis showed that *ak2*-deficient embryos presented a specific lack of both the dTOM and EGFP signals in LII.1 neuromasts from ~3 dpf on. Although the vast majority of the embryos analyzed did not present any positive signal in the secondary neuromasts at 5 dpf, surprisingly, in sporadic animals we observed dTOM^+^ cells between L1 and L2 neuromasts (Figure 3D, white arrowhead). Notably, at the same stage we never observed more mature hair cells (marked by *pou4f3*:GAP-GFP^+^) and, more importantly, these rare *atoh1a*:dTOM^+^ cells were never surrounded by *tnks1bp1*:EGFP^+^ supporting cells suggesting a depletion of the precursor cell pool used to populate the secondary neuromasts. This is similar to the depletion of hematopoietic stem and precursors cell observed in the zebrafish *ak2* alleles(*8*).

These data indicate that *ak2*-deficient embryos present an altered expression of markers for different neuromast cell populations, with a reduction in both immature and mature hair cell- and supporting cell-specific markers. However, the sporadic presence of solitary *atoh1a*:dTOM^+^ cells (Figure 3D, white arrow), in addition to the persistence in WISH analysis of a faint signal for the *eya1* marker in secondary neuromasts at 5 dpf (Figure 3C), suggest possible residual traces of secondary neuromasts in that region.

### *ak2* deficiency affects the ventral migration of secondary neuromasts and the development of interneuromast cells

To further verify the presence of residual secondary neuromasts between L1 and L2, we crossed *ak2^hg15/+^* animals with the transgenic reporter line Tg(−8.0*cldnb*:LY-EGFP). Until 4 dpf stage, this transgenic line expresses the EGFP reporter in all the different components of the PLL(*40*). From ~4 dpf, the EGFP expression in the PLL becomes specifically limited to the supporting cells in all neuromasts, the interneuromast cells and the afferent lateral line neurons innervating each neuromast(*41*, *42*). Confocal analysis of *ak2^hg15^* embryos and their siblings from 3 to 5 dpf showed that at 3 dpf some of the *ak2*-deficient embryos completely lacked EGFP signal in the LII.1 secondary neuromasts (Figure 4A). However, in the remaining *ak2^hg15^* population where the LII.1 retains some EGFP positive signal, secondary neuromast development was severely affected. In particular, we never observed the formation of EGFP^+^ LII.2 neuromasts. In addition, while wild-type LII.1 neuromasts migrate ventrally a few cell thicknesses, LII.1 neuromasts in *ak2* deficient larvae failed to migrate. In contrast to wild-type or carrier siblings, the neuromasts remained close to the horizontal myoseptum, at the same level of primary neuromasts (Figure 4A). Similar migratory defects in secondary neuromasts have only been previously described in the *strauss* mutants(*41*). Notably, at 5 dpf there is still a mixture of phenotypes among the *ak2^hg15^* embryos some that are totally negative for EGFP^+^ LII.1 neuromasts and some that possess the first secondary neuromast but no more. To confirm that the observed phenotypes were not due to early defects of the secondary neuromasts or the inability of primII to migrate, we also performed confocal analysis on *ak2^hg15^* Tg(−8.0*cldnb*:LY-EGFP) mutants and their siblings from 3.5 to 4 dpf (Video 1 and 2, respectively and Figure 4B). As summarized in Figure 4B, we did not observe major differences during primII migration between ~3 and 4 dpf.

**Figure 4.**
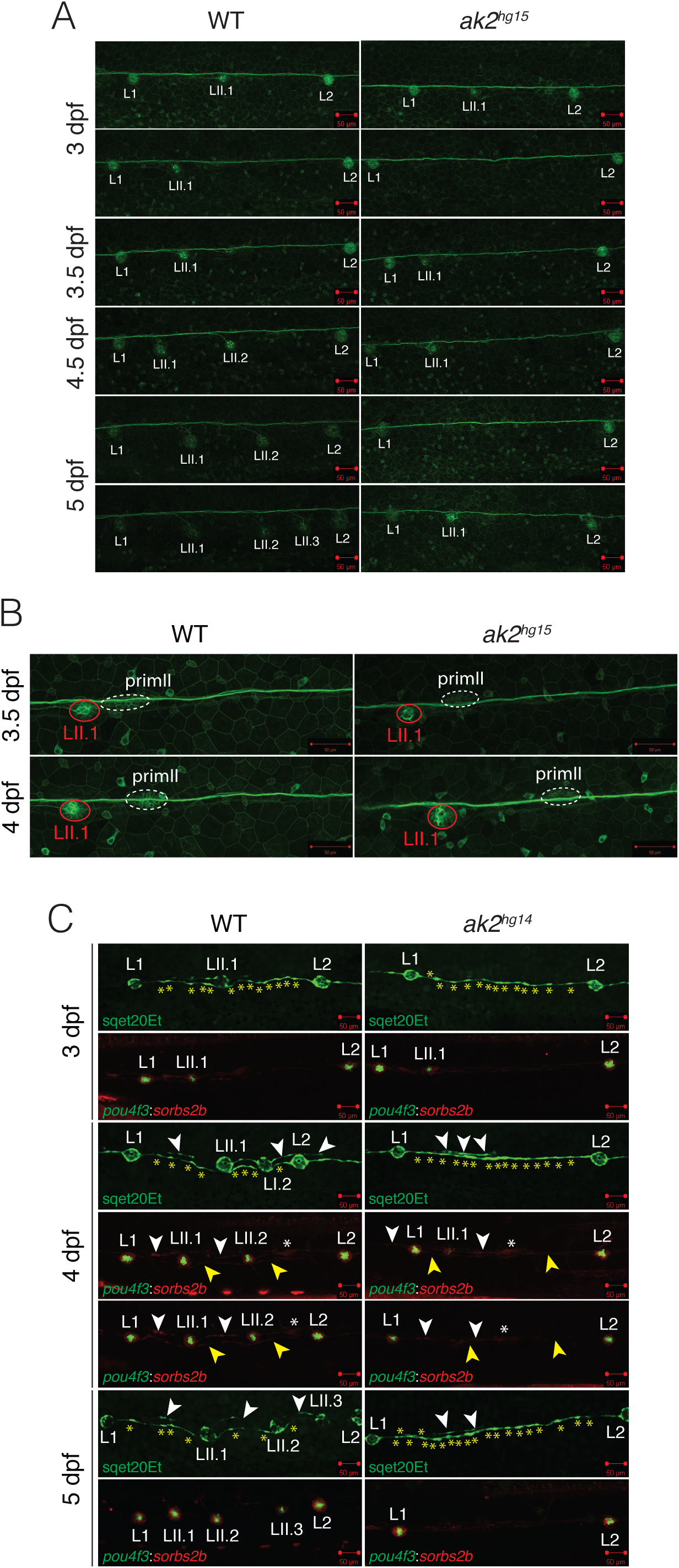
Analysis of lateral line phenotypes using different *ak2* mutant transgenic lines. (A) Confocal analysis at different developmental stages of the trunk regions of *ak2^hg15^* embryos and their control siblings in the Tg(*-8.0cldnb*:LY-EGFP) background labelling the migrating primordium, deposited neuromasts, and epithelial cells. (B) Confocal analysis from 3.5 to 4 dpf of trunk regions of an ak2*^hg15^* embryo and a control embryo (WT) in the Tg(-*8.0cldnb*:LY-EGFP) background labelling the migrating secondary primordium (dashed elliptic), deposited secondary neuromast (red circle), connecting interneuromast cells and epithelial cells. (C) Confocal analysis from 3 to 5 dpf of trunk regions of *ak2^hg14^* embryos and their siblings in different lateral line transgenic backgrounds. The sqet20Et line labels the mantle cells of rosettes as well as the interneuromast cells (green); *pou4f3*:*sorbs2b* indicates the Tg(*pou4f3*:GAP-GFP; mn0002Gt) double transgenic line labelling mature HCs (green) and supporting, mantle and interneuromast cells (red), respectively. Yellow asterisks label the interneuromast cells between L1 and L2 primary neuromasts in the sqet20Et transgenic line. White and yellow arrowheads denote primI or primII interneuromast cells, respectively. White asterisks label the migrating secondary primordium. Scale bars: 50 µm. L1 and L2 define primary neuromasts; LII.1, LII.2 and LII.3 designate the lateral line secondary neuromasts; primII indicates secondary primordium.

As previously mentioned, intercalary neuromasts derive from the proliferation and differentiation of the INCs, a population of precursor cells connecting the neuromasts of the lateral line that was deposited by the primI during its migration(*43*). In physiological conditions, this population of cells remains quiescent because of the presence of inhibitory signals deriving from glial cells(*44*, *45*), and isolated IC neuromasts can only rarely be observed between L1 and L2 primary neuromasts in WT embryos(*24*, *42*). However, it has been shown that, the lack of inhibitory signals induced by loss of glial cells and/or the PLL nerve(*44*, *45*) can prompt an ectopic production of IC neuromasts(*42*) in the region. Moreover, a previous study also showed that the ablation of the secondary primordium primII induces the ectopic production of IC neuromasts in the same region(*24*). Notably, in *ak2*-deficient embryos we never observed sporadic IC neuromasts nor an increased number of IC neuromasts between L1 and L2 neuromasts (Figures 2A, 3A-D and 4A), suggesting that the PLL glial and nervous cells are still present and secreting the inhibitory factors. However, the observed phenotypes could be that an *ak2* deficiency disrupts the formation of the interneuromast cells and IC neuromasts. Alternatively, it is formally possible the secondary neuromasts could still be present but lacking in the expression of the tested markers. To investigate these different possibilities, we performed confocal analysis from 3 to 5 dpf stages on two different transgenic lines in the *ak2^hg14/+^* background: the ET20 line labels mantle cells and interneuromast cells(*46*), and the double transgenic line Tg(*pou4f3*:GAP-GFP;GBT0002^mn0002GT^) with GFP expressed in hair cells and mRFP1 protein tagging the *sorbs2b* gene is expressed in supporting cells(*47*, *48*)(Figure 4C). As previously observed with other markers, the analysis showed an absence of mantle cells in the LII.1 secondary neuromasts (GFP negative), without apparent defects in the mantle cells of the primary neuromasts. We did occasionally observe interneuromast cells derived from primII migration (Figure 4C, white arrowheads) in *ak2^hg14^* embryos. Notably, from 4 dpf we detected an increased number of interneuromast cells between the L1 and L2 neuromasts (yellow asterisks), suggesting some active proliferation of the interneuromast cells derived from primI. In the *ak2^hg14^* mutants, primI- and primII-derived interneuromast cells and the primII primordium (yellow and white arrowheads and white asterisk, respectively) were mRFP1^+^ until 4 dpf. Moreover, only some of the null embryos had LII.1 neuromasts positive for hair and supporting cell markers, and it was at a very low level of expression (Figure 4C). Finally, at 5 dpf *ak2*-deficient embryos lacked the expression of both GFP and mRFP1 reporters between L1 and L2 primary neuromasts (Figure 4C) suggesting the cells are not viable long term.

Overall, these data suggest a possible defect in the activation of the interneuromast cells with a resulting deficiency in the formation of IC neuromasts. The mRFP1 expression in PLL secondary structures confirmed the presence of LII.1 secondary neuromasts in some of *ak2*-deficient embryos at least until 4 dpf but not after. These data are consistent with a premature depletion of the stem cell pool for the lateral line organs.

### Partial impairment of Wnt/β-catenin and Fgf signaling in *ak2*-deficient mutants

To verify that in *ak2*-deficient mutants the Schwann cells of PLL did not possess any severe deficiencies, we tested the effects of pharmacological inhibition of ErbB signaling using the tyrosine kinase inhibitor AG1478(*49*) on *ak2^hg14^* mutant and control embryos in the double transgenic background Tg(*tnks1bp1*:EGFP; *atoh1a*:dTOM). In particular, we focused our attention on the temporal window from 48-50 hpf, which corresponds to the end of the migration of the Schwann cells(*42*). Untreated and DMSO-treated *ak2^hg14^* embryos never produced *atoh1a*:dTOM^+^ and *tnks1bp1*:EGFP^+^ IC neuromasts (Figures 3D and 5A). As was previously reported(*42*), the AG1478 inhibition of ErbB signaling at 50 hpf was sufficient to induce the ectopic formation of IC neuromasts in all the control embryos (Figure 5A, top right panel). However, in *ak2^hg14^* mutants the treatment showed a partial penetrance, with three major classes of embryos: a) totally unresponsive (bottom right panel), b) partially responsive (middle right panel) and c) embryos possessing a number of ectopic neuromasts similar to controls (top right panel). Because at the tested dose we always observed a uniform response in the control population, the partial penetrance observed in the null mutants seems to reflect a real heterogeneity of the *ak2^hg14^* population’s response to the treatment and not just variation in inhibition levels. It is important to note that the induced neuromasts (white asterisks) were *atoh1a*:dTOM and *tnks1bp1*:EGFP double positive, although their size was strongly reduced compared to control siblings. Because we never observed the expression of these two markers in the secondary neuromasts of *ak2* mutants, their presence in AG1478-induced mutant neuromasts suggests a difference in the potency of secondary and IC neuromast precursors, probably due to their separate cellular origins.

**Figure 5.**
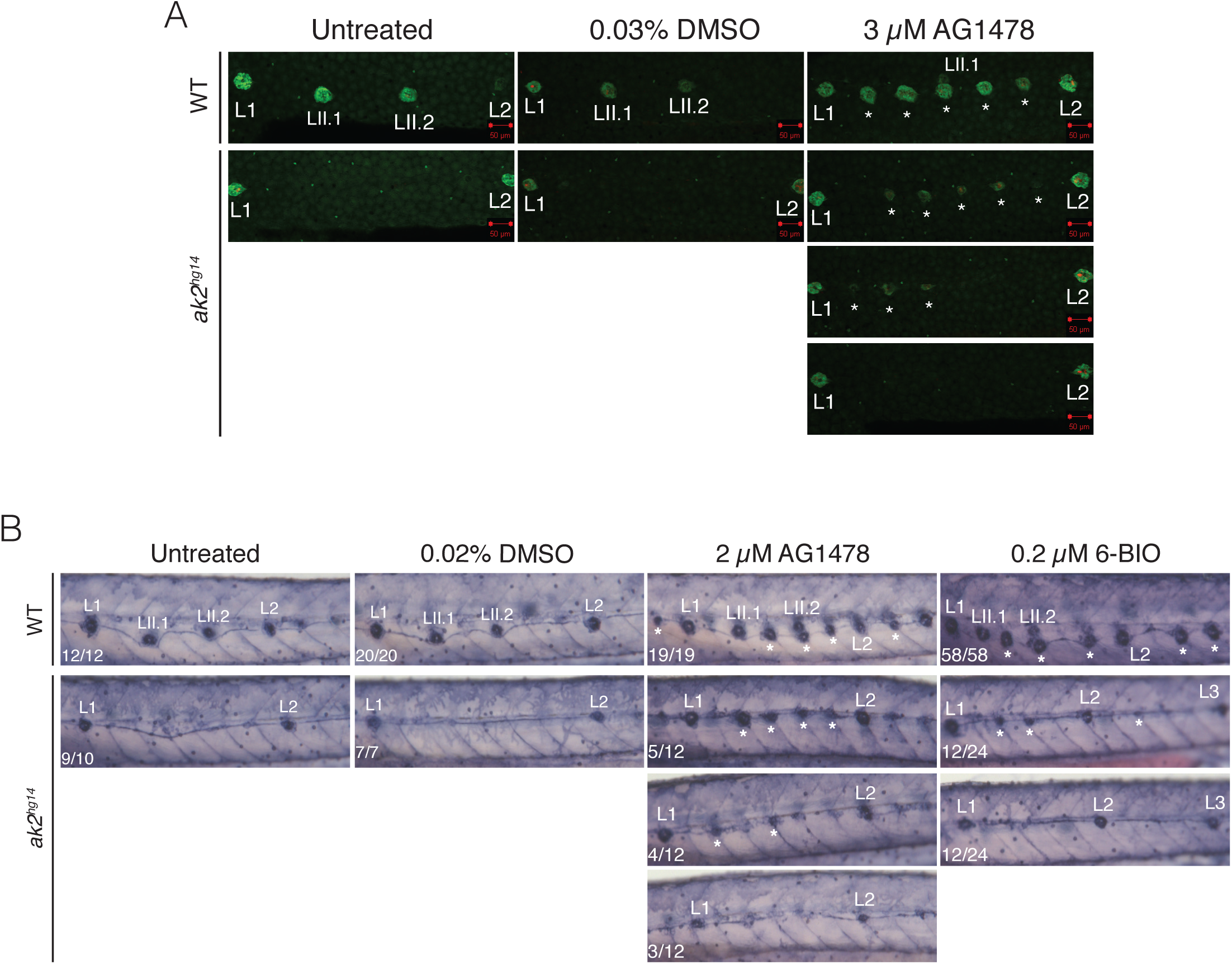
*ak2* deficiency impairs intercalary neuromast formation. (A-B) Chemical treatments of *ak2^hg14^* mutants and their control siblings with the ErbB tyrosine kinase inhibitor AG1478 or with 6-BIO, an activator of the Wnt signaling pathway through the inhibition of GSK-3. (A) Confocal analysis at 5 dpf of trunk regions for untreated, 0.03% DMSO or 3 µM AG1478 treated embryos in the Tg(*tnks1bp1*:EGFP; *atoh1a*:dTOM) transgenic background to visualize immature HCs (red) and supporting cells (green) of the neuromasts of the posterior lateral line. Scale bars: 50 µm. (B) Alkaline phosphatase staining to visualize the neuromasts of posterior lateral line of 5 dpf *ak2^hg14^* and control embryos untreated or treated with 0.02% DMSO, 2 µM AG1478 or 0.2 µM 6-BIO compounds. For each specific phenotype, the number of embryos observed over the total number of analyzed embryos are indicated on the bottom left corner. L1 and L2 define primary neuromasts; LII.1 and LII.2 designate the lateral line secondary neuromasts; the asterisks indicate intercalary neuromasts.

Although little is known about the signaling pathways that orchestrate IC neuromast formation, a previous study showed that it requires both Wnt/β-catenin and Fgf signaling pathways(*42*). In particular, while the Wnt/β-catenin signaling is required for initial interneuromast proliferation, Fgf signaling is crucial for subsequent organization of interneuromast cells into a rosette structure and for triggering cellular differentiation(*42*). To determine if these pathways were affected in *ak2* mutants, we treated the embryos from an *ak2^hg14/+^* incross with 6-BIO, a potent inhibitor of the GSK-3 enzyme (Figure 5B). To obtain a quantitative analysis, we performed an AP staining of the embryos and we included an AG1478 treatment as further internal control. 6-BIO has been shown to result in the strong activation of the Wnt/β-catenin pathway(*50*) and the formation of extra IC neuromasts in the PLL of treated zebrafish embryos(*42*). As shown in Figure 5B, in *ak2^hg14^* untreated or DMSO-treated embryos secondary or IC neuromasts between L1 and L2 were always AP negative. At the same time, 100% of the AG1478 and 6-BIO-treated control embryos developed extra IC neuromasts. It’s important to note that, in order to avoid 6-BIO toxicity from 24 hours of administration, we used it at a concentration of 0.2 µM (which is 10 times lower than the previously published dose(*42*, *51*)). After 3 days of treatment with the selected dose, we observed a strong induction of ectopic neuromasts in the trunk regions of control embryos, without any major morphological defects or reduced survival of the embryos. These results were very similar to those we obtained on control embryos with a 2 µM AG1418 treatment (Figure 5B). As previously observed (Figure 5A), the *ak2*-deficient embryos treated with the AG1478 compound could be classified in three different groups. The AG1478 administration induced the formation of ectopic intercalary neuromasts in ~42% (5/12), a partial response in ~33% (4/12), and a lack of intercalary neuromast formation in the remaining *ak2*-deficient population (25%, 3/12). Similarly, we found a partial penetrance using the GSK-3 inhibitor 6-BIO, as well. However, in this case we observed that half of the *ak2*-deficient population (12/24) was totally unresponsive to the treatment, while the other half showed a very mild response, with a reduced number of ectopic IC neuromasts compare to controls. In both the treatments we noticed that, again, the ectopic IC neuromasts were significantly reduced in their size (Figure 5A-B).

Overall these data suggest an impairment of the Wnt/β-catenin and FGF signaling pathways in the interneuromast cells of *ak2*-deficient embryos. The increased proliferation observed in the interneuromast cells of *ak2* null mutants (Figure 4C) suggest that, in basal conditions, the interneuromast cells seem to be able to initially respond to the Wnt/β-catenin pathway. However, without an external stimulus provided (like AG1478 or 6-BIO treatments which induce a sustained Wnt/β-catenin activation), interneuromast cells fail to generate ectopic intercalary neuromasts. Because it has been shown that the Wnt/β-catenin pathway is upstream of FGF, and that only Fgf signaling is required for subsequent rosette formation, the consistently reduced number and size of rosettes observed in *ak2-*deficient mutants after AG1478 or 6-BIO treatments suggest a partial impairment of the FGF signaling as well.

### *ak2* deficiency induces increased cell death in inner ear, primII and PLL neuromasts

Previous studies showed that, in RD patient fibroblasts and zebrafish hematopoietic tissues, ak2 protein can possess an antiapoptotic function(*2*, *8*). Therefore, we wondered if the same phenomena could be the cause of the decreased number of hair cells observed in *ak2*-deficient animals.

To investigate the possible presence of dying hair cells in the inner ear and the PLL structures, we performed TUNEL assays on *ak2*-deficient embryos and their siblings in the *pou4f3*:GAP-GFP or *cldnB*:LY-EGFP transgenic backgrounds, respectively. Because the organs of interest were on the larval skin, we had to use an optimized TUNEL protocol with a very mild digestion of the embryos (see the Material and Methods section for further details). In general, TUNEL staining from 2 to 4 dpf showed an increased level of apoptotic cells in *ak2^hg15^* embryos compared to the controls (red signal in Figure 6, Figure 6-figure supplement 1 and data not shown). The analysis of the TUNEL signal distribution in the area surrounding the inner ear in each section of the whole Z-stack allowed us to verify that, in most cases, the TUNEL positivity corresponded to the nuclei of mature *pou4f3*:GAP-GFP^+^ cells in the neuromasts of the anterior lateral line and in the inner ear structures (yellow arrowheads in Figure 6-figure supplement 1).

**Figure 6.**
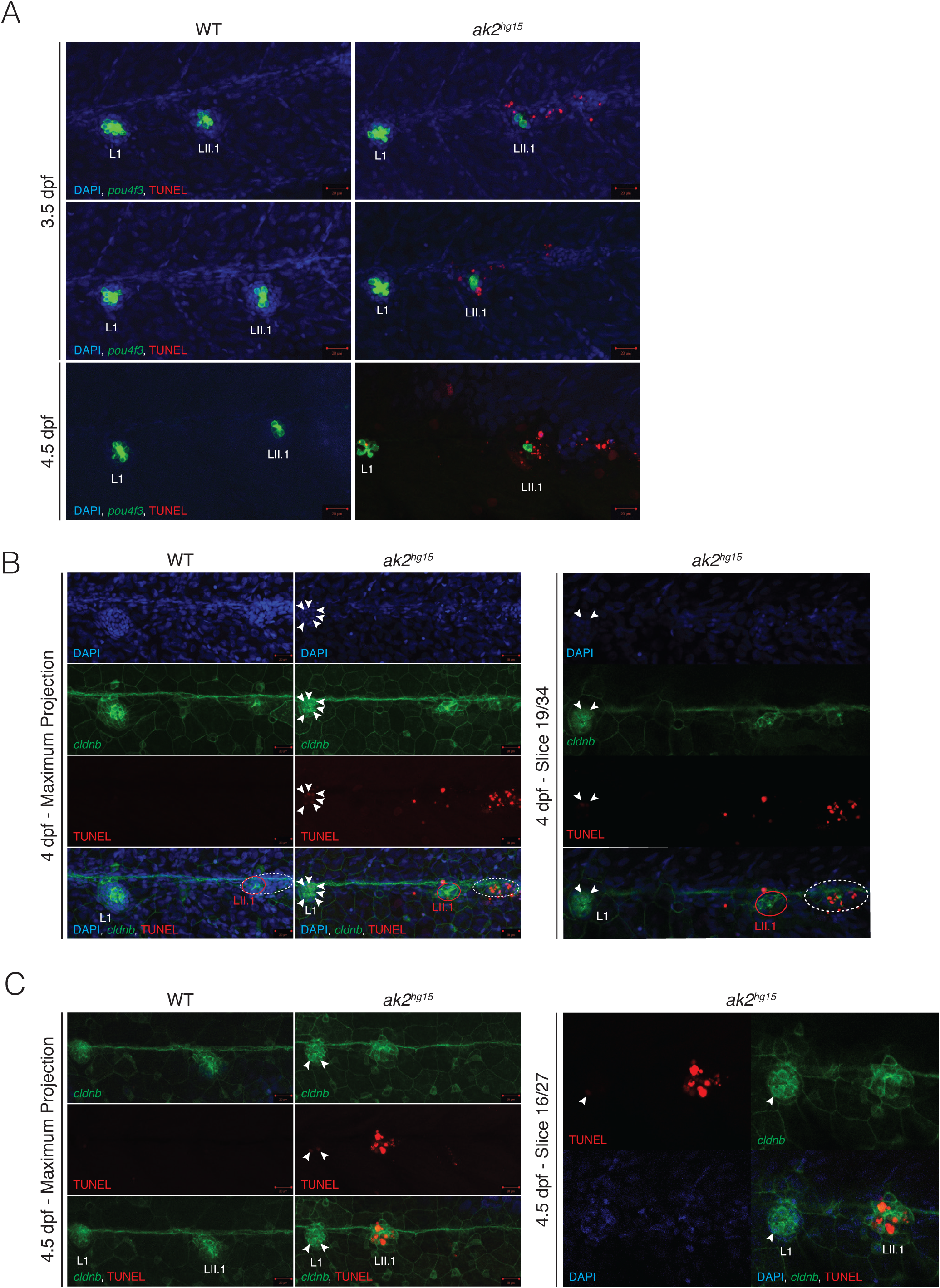
Increased cell death in the otic vescicles and the posterior lateral line of *ak2^hg15^* embryos. (A) Maximum projections of 3.5 and 4.5 dpf ak2*^hg15^* embryos and their control siblings in the Tg(*pou4f3*:GAP-GFP) background (indicated as *pou4f3*) stained with TUNEL assays (red signal). (B-C) Maximum projections (left panels) and representative single plane confocal analysis (right panels) at 4 (B) and 4.5 (C) dpf of a TUNEL assay (red signal) performed on *ak2^hg15^* embryos and their siblings in the Tg(-*8.0cldnb*:LY-EGFP) background (*cldnb*, green) labelling the migrating secondary primordium (dashed elliptic), deposited primary and secondary (red circle) neuromasts and epithelial cells. White arrows label TUNEL positive cells in the L1 primary neuromast. Nuclei are labelled with DAPI (blue). L1 and L2 are primary neuromasts; LII.1 designates the lateral line secondary neuromast. Scale bars: 20 µM.

Notably, for the same samples we also observed a strong signal in the trunk regions between L1 and L2 primary neuromasts, suggesting the presence of apoptotic cells in structures that could represent the primII and the secondary neuromasts (Figure 6A). To verify this possibility, we repeated the analysis on 4 dpf trunk regions of *ak2* mutants in the *cldnb*:LY-EGFP transgenic background. As shown in Figure 6B, the analysis confirmed the presence of a striking number of TUNEL^+^ cells in the migrating primII, with very few dead cells in the secondary neuromast LII.1. At the same stage we observed the presence of several TUNEL^+^ cells in L1 primary neuromasts (white arrowheads in Figure 6B), providing an explanation for the reduced number of hair cells (in combination with a regeneration deficiency) in primary neuromasts previously observed using Yo-PRO-1 staining or transgenic backgrounds (Figures 2C-D and Figure 2-figure supplement 2). We obtained similar results in LII.1 secondary neuromasts at 4.5 dpf (Figure 6C) which correlated with the strong reduction of *eya1* observed at similar stages (Figure 3C).

It appears that the *ak2* deficiency induced increased levels of cell death in zebrafish mechanosensory organs combined with an inability to regenerate properly, explaining the reduced number of mature hair cells in the inner ear and the developmental and regenerative defects observed in primary and secondary neuromasts.

### *ak2*-deficiency triggers oxidative stress response genes

Ak2 deficiency has been linked to an increased level of intracellular ROS and oxidative stress(*2*, *8*). In particular, in the hematopoietic tissues of the zebrafish *ak2* models of RD we previously described the induction of heme oxygenase 1a (*hmox1a)* expression (*8*), which represented a mechanism for cells to defend against potential damage caused by increased oxidative stress(*52*, *53*). The expression of *hmox1a* and other antioxidant genes is regulated by the binding of the Nrf2 transcription factor to specific enhancer elements known as AREs (or antioxidant response elements)(*52*, *54*). In order to characterize the possible effects of increased levels of oxidative stress induced by the *ak2* deficiency in sensory organs, we tested the expression of several genes involved in the cellular defense against ROS, including some specific *nrf2* target genes(*54*–*56*) (Figure 7 and Figure 7-figure supplement 1). Some of these markers were expressed at detectable levels in the inner ears (*prdx4-6*, *gpx1a-b* and *gpx4a-b*) and PLL neuromasts (*prdx2* and *gstp1)* under normal physiologic conditions, indicating that they normally participate in a basal antioxidant defense in these structures. Notably, in *ak2*-deficient embryos we did not observe a uniform increase of expression across the whole set of the markers, as initially predicted based on the presumed high levels of oxidative stress in mutant embryos. Based on how the stress genes responded (up, down or unchanged expression) in the inner ear or the PLL, we divided the genes in three categories (Figure 7 and Figure 7-figure supplement 1). Figure 7A shows that most of the peroxiredoxin (*prdx*) genes showed an increased level of expression in the inner ear. Notably, *prdx1* expression is also upregulated in several other tissues of the body, including the PLL (Figure 7A and data not shown). Among the *prdx* gene family, the only exception was *prdx2*, which was not expressed in the inner ear and whose expression was reduced in caudal hematopoietic tissue (CHT) and in the primary and secondary neuromasts of the PLL (Figure 7C). Other examples of genes that were down-regulated in *ak2*-deficient embryos were the glutathione peroxidase 4 genes (*gpx4a* and *gpx4b*); in particular the *gpx4b* expression appeared strongly reduced in the inner ears (Figure 7B). The expression of glutathione peroxidase 1a and 1b (*gpx1a* and *gpx1b*) and superoxide dismutase 1 and 2 (*sod1* and *sod2*) appeared to be unchanged between *ak2*-deficient embryos and their control siblings (Figures 7-figure supplement 1A and 1B, respectively).

**Figure 7.**
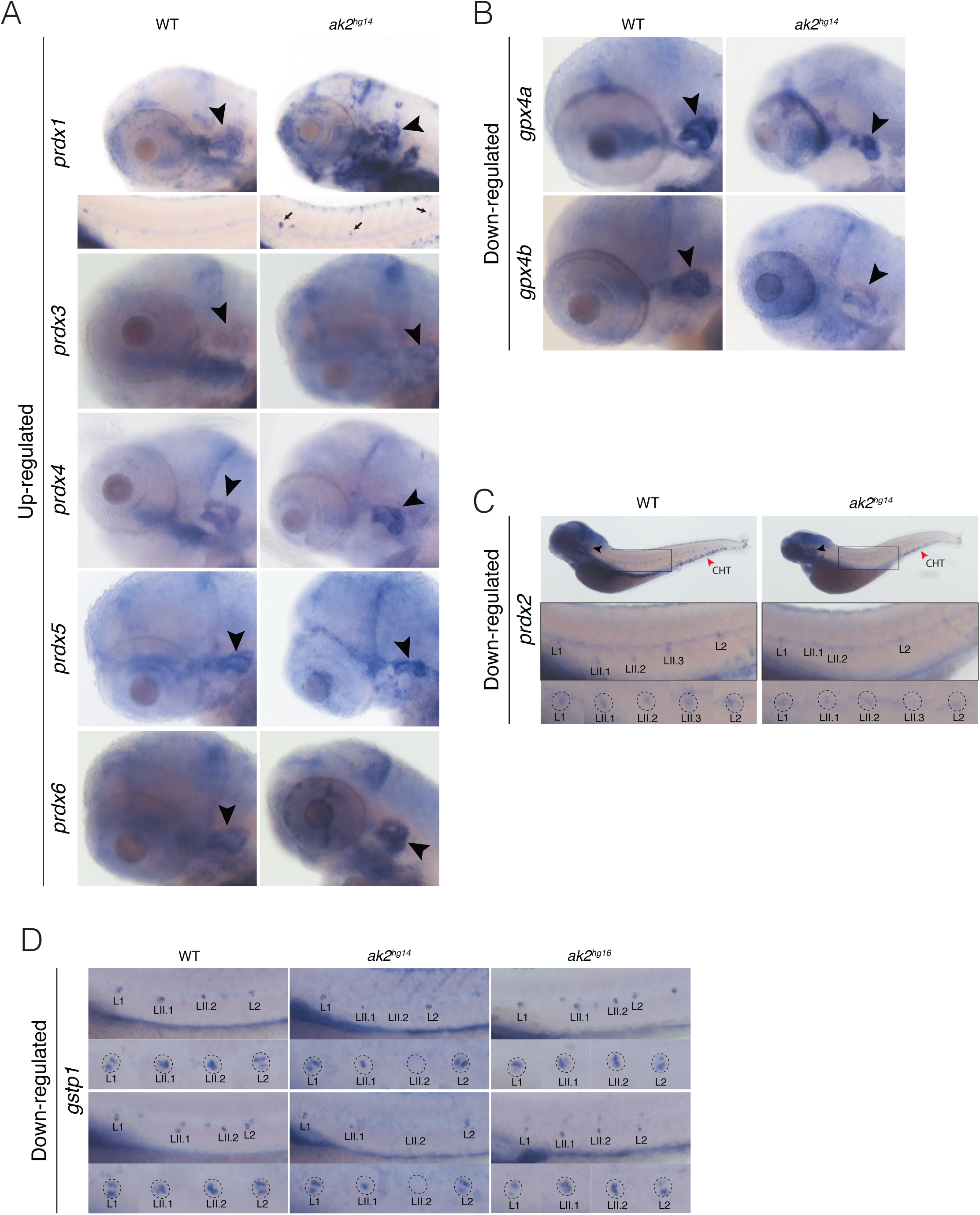
Altered expression of oxidative stress markers in *ak2^hg14^* embryos. Expression analysis of oxidative stress markers in *ak2^hg14^* and control sibling embryos at 4 or 5 dpf. Lateral views with anterior to the left. Black arrowheads indicate the otic vescicle. Markers were grouped in three different classes based on the expression level of each gene in otic vesicle and/or PLL in nulls and control embryos. (A) Up-regulated expression of several *prdx* genes in the otic vesicle and posterior lateral line at 4dpf. Bottom panel for prdx1 marker: the black arrows indicate the ectopic expression of the marker in PLL neuromasts of *ak2^hg14^* embryos. (B) Down-regulated expression of *gpx4a* and *gpx4b* markers at 4 dpf. (C) Specific down-regulation of *prdx2* marker at 5 dpf in posterior lateral line neuromasts (dashed circles in specific insets of black boxes) and caudal hematopoietic tissue (red arrowhead, CHT). (D) Comparison of *gstp1* expression in the posterior lateral line neuromasts in control embryos, *ak2^hg14^* null and *ak2^hg16^* hypomorphic mutants at 4 dpf. Two different embryos per genotype are shown. L1 and L2 define primary neuromasts; LII.1, LII.2 and LII.3 designate the lateral line secondary neuromasts. *gpx*: glutathione peroxidase; *gstp1*: glutathione S-transferase pi 1; *prdx*: peroxiredoxin; *sod*: superoxide dismutase.

Surprisingly, at 4 dpf the pattern of expression of glutathione S-transferase pi 1 (*gstp1*) gene seemed to differ significantly between the primary and secondary neuromasts (Figure 7D). In the secondary LII.2 neuromasts, which are deposited later than LII.1 neuromasts, *gstp1^+^* cells were typically limited to the central part of the neuromast, likely representing the most mature hair cells of the neuromast. Older primary neuromasts like L1 and L2 presented a specific and repeatable pattern of expression, with *gstp1^+^* cells mostly limited to the dorso-ventral poles of the neuromast and, usually, with an asymmetrical level of expression between the two poles. In the central part of the more mature neuromasts, *gstp1* was expressed at a very low level or not at all (Figure 7D). Secondary LII.1 appeared to represent an intermediate state between the older neuromasts and LII.2, with a broader number of *gstp1^+^* cells and with one pole, in this case on the antero-posterior axis, showing higher expression of the marker. This difference in the axis orientation correlated with the different hair cell polarity between primary and secondary neuromasts, which possess an antero-posterior or dorso-ventral orientation, respectively(*24*, *41*). The variation in the *gstp1* expression pattern seemed to correlate with the different maturation state of each neuromast, consistent with the *gstp1* expression data at earlier stages (2-3 dpf) shown in Figure 7-figure supplement 1C and those publicly available through ZFIN indicating a clear expression of the gene in the most central part of the primary neuromasts (*57*, *58*). Compared to control siblings or *ak2^hg16^* hypomorphic mutants, *ak2*-deficient embryos completely lacked *gstp1* expression in the LII.2 neuromast; while in LII.1 neuromasts its expression was reduced and restricted only to the most central cells of the neuromast. These data were supported by our previous observations using Yo-PRO-1 staining and confocal analysis (Figure 2B, 3D and 4C), suggesting that *ak2*-deficiency impairs the maturation of the LII.1 neuromasts.

Taken as a whole, these results confirm that an *ak2* deficiency perturbs the physiologic oxidative state of the cells inducing an altered expression of some but not all oxidative stress-related genes. The altered expression of zebrafish NRF2 targets such as *prdx1* and *gstp1* were not uniformly increased suggesting the specific cellular context could alter the responses of different stress markers and potentially could explain how different tissues displayed different pathologies.

### Antioxidant treatment partially rescued inner ear hair cell and PLL neuromast defects

Previously, we showed that antioxidant treatment was able to partially rescue the hematopoietic phenotypes observed in the zebrafish models of RD(*8*). In particular, a treatment with 10-50 µM glutathione reduced the ectopic expression of the oxidative stress marker *hmox1a* and also restored the expression of *c-myb* and *rag1* (markers of hematopoietic stem cells and lymphoid populations, respectively) in the hematopoietic tissues of *ak2*-deficient embryos(*8*). In addition, the treatment of RD patient iPSCs with glutathione (GSH) was able to increase the *in vitro* differentiation rate of neutrophils(*8*). We therefore tested if a similar treatment would be beneficial to the sensorineural defects of a zebrafish RD model. We treated the *ak2^hg14^* and control embryos with a range of GSH concentrations. Using the *pou4f3*:GAP-GFP transgenic background, we initially compared the number of mature hair cells in the inner ear cristae and the anterior macula of 3 dpf *ak2^hg14^* and control embryos, untreated or treated with 100-300 µM GSH (Figure 8-figure supplements 1); then we extended the analysis to the cristae of 4 dpf *ak2^hg14^* and control embryos in the same conditions (Figure 8-figure supplements 2). The comparison of the number of hair cells in control and mutant animals in different conditions (untreated vs different doses of the treatment) allowed us to calculate the % of rescued hair cells induced by the treatments (Figure 8C). At any of the tested doses the treatment significantly affected the number of hair cells of control embryos with an average of ~10, ~28 and ~20 hair cells in the 3 dpf cristae and macula, and in 4 dpf cristae, respectively (Figure 8-figure supplements 1 and 2). The specific comparison between untreated and GSH-treated *ak2^hg14^* mutants at 3 dpf showed a statistically significant increase in the average number of hair cells in both the cristae and the anterior macula of null mutants (Figure 8A, top and bottom panels, respectively). In particular, we observed an average of 6.9, 6.8 and 7.4 hair cells in the cristae using increasing doses of GSH which correspond to 17%, 16% and 22% of rescued hair cells, respectively (Figure 8C). Similar results were also obtained in the cristae of 4 dpf *ak2^hg14^* and control embryos (Figure 8B and Figure 8-figure supplement 2), where we observed a dose dependent response for the rescue that was statistically significant, and the antioxidant treatments were able to rescue the 16%, 25% and 33% of the hair cells, respectively (Figure 8B and 8C).

**Figure 8.**
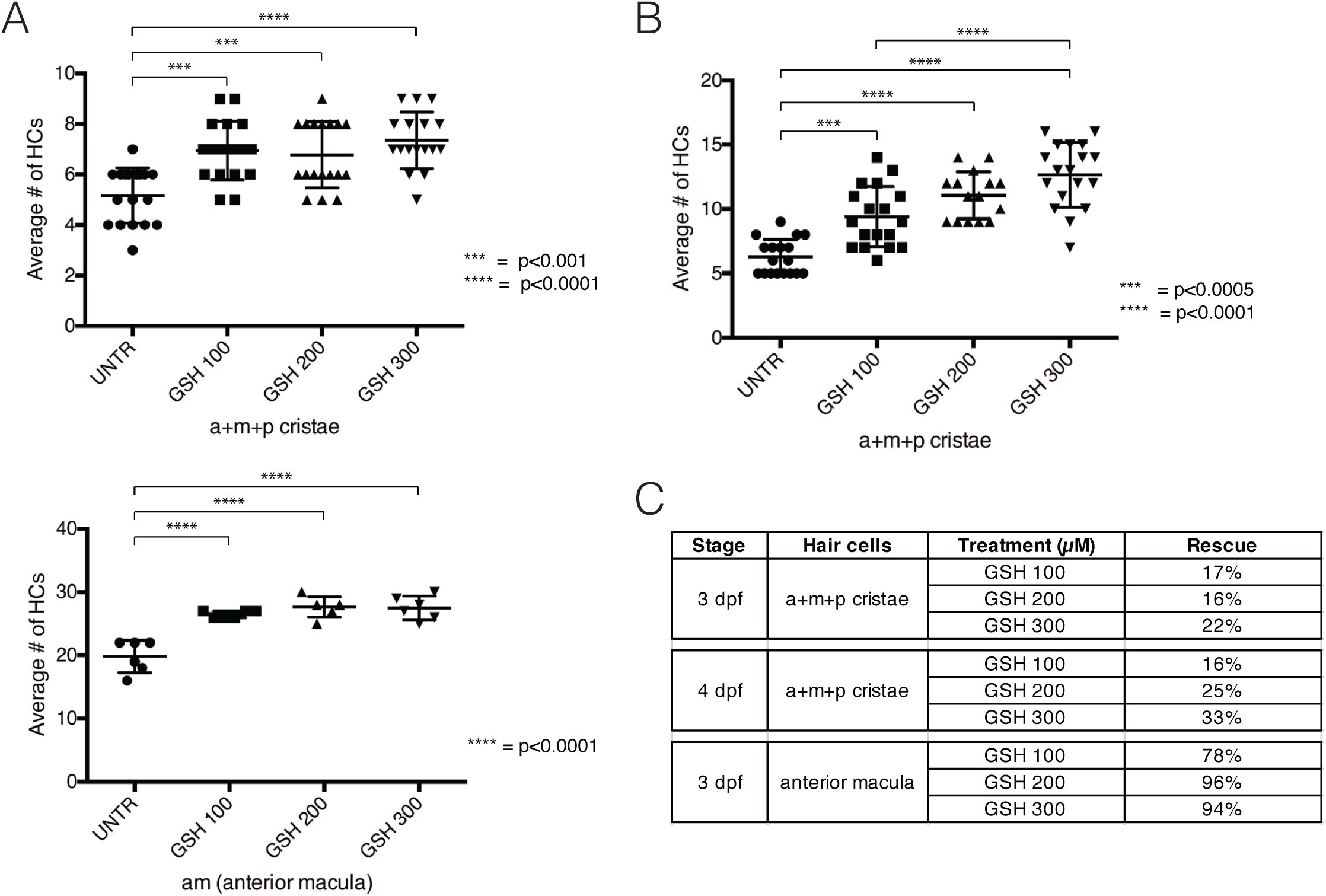
Antioxidant treatment partially rescues HC numbers in inner ear of *ak2^hg14^* mutants. (A) Comparison of the average number of mature hair cells in the inner ear cristae (upper graph) or anterior macula (bottom graph) of 3 dpf *ak2^hg14^* embryos treated with different concentrations of GSH (100-300 µM). UNTR: untreated. (B) Comparison of the average number of mature HCs in the inner ear cristae of 4 dpf *ak2^hg14^* embryos treated with different concentrations of GSH (100-300 µM). UNTR: untreated. (C) Summary of the % of rescue induced by the different GSH treatments on the inner ear cristae and anterior macula at 3 and 4 dpf. (A-B) A one-way ANOVA analysis (followed by Tukey’s post hoc test; p<0.001) was used to compare groups of fish treated differently. Results are shown as mean +/− standard error of the mean. a: anterior; m: medial; p: posterior.

Finally, the analysis of the average number of hair cells in the 3 dpf macula of *ak2^hg14^* mutants and control siblings showed that the macula seems to be less sensitive to the *ak2* deficiency with an average of ~20 hair cells (~28 in the controls) which correspond to a 29% of reduction of hair cells, compared to the 48% observed in the cristae of untreated embryos at the same stage or the 69% at 4 dpf. Notably, the GSH treatments were able to almost fully restore the physiological number of hair cells in this specific structure with a 78%, 96% and 94% of rescue. This may indicate that the antioxidant agent might be more effective during the first phases of the *ak2* deficiency, suggesting that they might work best when the damage at the target tissue is of a reduced extent. Unfortunately, because of the anatomical localization of the macula and the consequent difficulties to obtain a good imaging of the structure, we were not able to repeat the analysis at 4 dpf or later, to see if the strong recovery observed at 3 dpf is sustained during time.

We also tested the ability of a GSH treatment to rescue secondary neuromast development. As previously shown, from ~4 dpf secondary neuromasts in *ak2^hg14^* mutants were *atoh1a*, *atoh1a*:dTOM, *tnks1bp1*:EGFP and *pou4f3*:GAP-GFP negative (Figures 3A, 3D, 4C, 5A and 9A). At 5 dpf they usually were also *cldnb*:LY-EGFP negative and they partially or totally lacked alkaline phosphatase expression (Figure 2A, 4A, 5B and 9B). When we treated the embryos with 100 µM GSH, we observed a partial rescue of the secondary neuromast phenotypes. In particular, we observed the recovery of *atoh1a*:dTOM and *tnks1bp1*:EGFP expression in secondary LII.1 neuromasts at 4 dpf (Figure 9A, white arrowheads). Although at 5 dpf the secondary neuromasts also showed *cldnb*:LY-EGFP expression (Figure 9A, yellow arrowheads), we never observed the recovery of *pou4f3*:GAP-GFP expression, suggesting that the antioxidant treatment may not be sufficient to induce the full maturation of the hair cells in secondary neuromasts. To perform a more quantitative analysis of the rescue induced by the GSH treatment in the embryos and to clearly distinguish secondary and intercalary neuromasts, we repeated the same analysis using two different doses of GSH (100 and 200 µM) and then performed an AP staining at 4 and 5 dpf. As shown in Figure 9B, at 4 dpf ~58% (7/12) of the untreated *ak2^hg14^* embryos had no expression of the AP in secondary neuromasts. However, the 100 µM and 200 µM GSH treatments dropped that percentage to 47% (8/17) and 33% (6/18), respectively. At 5 dpf, we did not observe any AP signal in secondary neuromasts in untreated embryos (Figure 9B, left column); however, we observed a partial rescue of AP activity in 3/14 (21%) and 4/11 (36%) embryos treated with 100µM and 200µM GSH, respectively.

**Figure 9.**
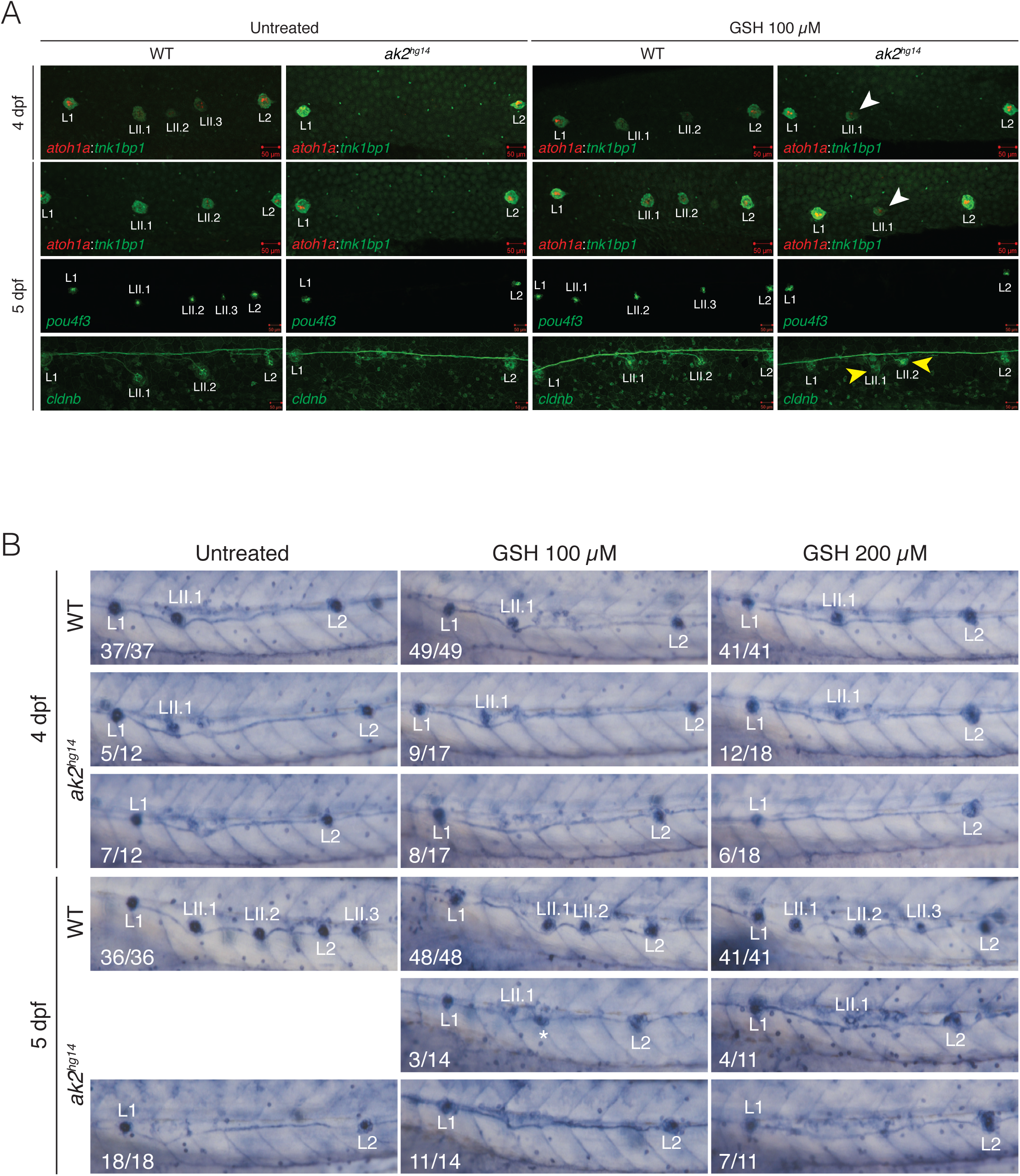
Antioxidant treatment partially rescues secondary neuromast defects in *ak2^hg14^* mutants. (A) Confocal analysis from 4 to 5 dpf of trunk regions of *ak2^hg14^* embryos and their siblings in different transgenic background to visualize different components of posterior lateral line. Untreated embryos are compared to GSH 100 µM treated embryos. *pou4f3* denotes the Tg(*pou4f3*:GAP-GFP) line labelling the mature hair cells (green); *atoh1a*:*tnks1bp1* indicates the Tg(*tnks1bp1*:EGFP; *atoh1a*:dTOM) line labelling immature hair cells (red) and supporting cells (green), respectively. *cldnb* denotes the Tg(-*8.0cldnb*:LY-EGFP) labeling all cells in the deposited neuromasts and the connecting interneuromast cells. White and yellow arrowheads indicate the rescued expression of specific transgenic markers in secondary neuromasts. Scale bars: 50 µm. (B) Alkaline phosphatase staining to visualize the neuromasts in the posterior lateral line of 4 and 5 dpf *ak2^hg14^* and control embryos untreated or treated with GSH (100 or 200 µM). For each specific phenotype, the number of embryos observed over the total of analyzed embryos are indicated on the bottom left corner. L1 and L2 label primary neuromasts; LII.1, LII.2 and LII.3 designate the lateral line secondary neuromasts.

Taken together, these data indicate that, as previously observed for hematopoietic phenotypes, the administration of an antioxidant treatment (in this specific case GSH) can partially reduce the sensorineural defects observed in an *ak2*-deficient zebrafish model of RD; confirming that these treatments could be potentially helpful in ameliorating a wide range of symptoms in RD patients.

## Discussion

Reticular dysgenesis is a rare and severe immunodeficiency also characterized by the presence of sensorineural deafness. In the present study, we took advantage of two different zebrafish models of the disease to extend our knowledge of the physiopathology of RD hearing loss, demonstrating that *ak2* is expressed in vertebrate sensory organs and that it has a crucial role in zebrafish sensory system development. The phenotypes observed in our models of *ak2* deficiency could explain the mechanism of action resulting in the sensorineural deafness of RD patients. Ak2 deficiency, through increased oxidative stress and cell death, reduces the total number of mature hair cells in the inner ear of mutant animals and it also impairs their regenerative ability. Moreover, it specifically affects the formation of primary and secondary neuromasts of the posterior lateral line. Finally, we proved that, as previously observed for hematopoietic defects, ameliorating the oxidative stress with an antioxidant treatment partially rescues the sensorineural phenotypes observed in the mutant larvae. However, the lower range of doses required to reduce the oxidative stress and to restore the expression of HSC and lymphoid markers in *ak2* null embryos suggest that hematopoietic cells seems to be more sensitive to the oxidative stress than cells from sensory system (and thus easier to rescue with antioxidants alone). This idea seems to be supported by the fact that we never observed sensorineural defects in the hypomorphic *ak2^hg16^* animals, although they present the same severe hematological phenotypes observed in *ak2*-deficient embryos(*8*).

It is interesting to note that the antioxidant treatments presented a different level of efficacy on the two different parts of the zebrafish sensory system. In the inner ear of *ak2*-deficient mutants the treatment was able to partially rescue the number of mature hair cells (*pou4f*3-GFP^+^) in the cristae and to almost fully restore their correct number in the macula. However, the effect antioxidants had on secondary neuromasts of the PLL was limited to the induction of some of the molecular markers of early neuromasts (*atoh1a*:dTOM and *tnks1bp1*:EGFP, *cldnb*:LY-GFP, and partially alkaline phosphatase) while we never observed the formation of more mature *pou4f3*:GAP-GFP*^+^* hair cells. Although there are many potential reasons for this observation, one possible explanation could be that it reflects a major difference in the expression profile of hair and supporting cells between primary and secondary neuromasts, with secondary neuromasts relying more heavily on *ak2* as opposed to other adenylate kinases encoded in the genome. Alternatively, it could also be due to a distinctive role of *ak2* during the different stages of neuromast maturation. In particular, our data seems to suggest that during earlier stages of neuromast development, as observed in secondary neuromasts, *ak2* could be involved in maintaining a correct transcriptional profile of the cells possibly via metabolic feedback. Eventually, during the later phases of neuromast development, *ak2* role could be more directly linked to the cell survival, as suggested by the increased level of apoptotic cells in inner ear and primary neuromasts. Notably, altered transcriptional profiles have been previously observed in two different *in vitro* models of RD(*10*, *11*). Further investigations will be required to test this hypothesis. Another possibility is that some pluripotent or multipotent stem cell populations are sensitive to ak2 deficiencies and are incapable of sustained self-renewal, eventually showing stem cell “exhaustion.”

A variety of mutations have been isolated in RD patients, ranging from missense and nonsense mutations affecting single amino acid positions or the AK2 pre-mRNA splicing, to a range of deletions(*1*, *2*, *4*, *5*, *28*, *59*–*61*). Although the relative impact from the different genomic mutations has not been systematically evaluated, in most of the cases the mutations appear to induce a strong reduction of the AK2 protein level, without significantly affecting the mRNA levels(*1*, *2*, *5*, *62*). A surprising finding with our RD models was that *ak2^hg16^* hypomorphic mutants were able to reach adulthood despite the severe hematological defects observed. This hematopoietic phenotype was shared with the *ak2^hg14^* and *ak2^hg15^* alleles which were lethal, leading us to believe the mortality in the stronger alleles was not linked to the initial hematopoietic defects (*8*). WISH analysis showed that *ak2^hg16^* mutants presented normal levels of *ak2* mRNA expression (although protein levels were not determined), while frame-shift mutations in exon 1 in the *ak2^hg14^* and *ak2^hg15^* alleles triggered nonsense-mediated decay of *ak2* mRNA(*8*). The lack of defects affecting the sensory systems both during larval and adult phases in *ak2^hg16^* hypomorphic mutants despite a strong hematopoietic phenotype strongly suggests that, at least in zebrafish, the hematopoietic lineages are significantly more sensitive to ak2 reduction than the sensorineural cells. So far, with only one exception(*28*), no other major non-hematological defects other than sensorineural hearing loss have been reported in RD patients(*5*) despite seeing several affected tissues in the zebrafish models of the disease. Identifying all the possible compromised tissues in RD patients will be essential to understanding why bone marrow transplants alone are not sufficient to completely cure the disease.

## Material and Methods

### Zebrafish Lines and maintenance

Zebrafish were maintained and used following protocols approved by the National Human Genome Research Institute Animal Care and Use Committee. All protocols and methods related to animals or animal tissues were approved by animal care and use committee of the National Human Genome Research Institute (G-01-3). Zebrafish handling, breeding, and staging were performed as previously described(*63*, *64*). To prevent pigmentation, the embryos used for WISH or confocal analysis were cultured in fish water containing 0.003% 1-phenyl-2-thiouera (PTU, Sigma-Aldrich, cat. # P7629) from 24hpf on. The following strains were used: TAB5(*65*); Tg(-*8.0cldnb*:LY-EGFP)(*40*) (also indicated in the text as *cldnb*); sqet20Et(*46*, *66*); Tg(*tnks1bp1*:EGFP;*atoh1a*:dTOM) (*39*); Tg(*pou4f3*:GAP-GFP)(*67*) (also designated in the text as *pou4f3*); and the Sorbs2b:mRFP1 insertional transgenic line(*68*) in Tg(*pou4f3*:GAP-GFP) background Tg(*pou4f3*:GAP-GFP; mn0002Gt) (designated in the text as *pou4f3:sorbs2b*).

### Whole mount in situ hybridization (WISH)

WISH analysis were performed as previously described(*8*, *69*), with minor modifications. For *ak2* antisense probes, the Hybridization Mix was supplemented with 5 % w/v Dextran Sulfate(*69*). The DIG RNA Labeling kit (SP6/T7; Roche) was used for *in vitro* transcription using PCR products cloned into the pCR4-TOPO vector (Life Technologies) or directly from PCR fragments(*69*). All probes were purified using Spin Post-Reaction Clean-Up Columns (Sigma-Aldrich) before use. The following DIG-labeled antisense mRNA probes were generated: *eya1*, *atoh1a*, *prdx1-6*, *gpx1a-b*, *gpx4a-b*, *sod1-2*, *gstp1*. See Table S1 for a complete list of primers. All embryos used for WISH were fixed overnight in 4% formaldehyde (PFA)/PBS, rinsed with PBS-Tween, dehydrated in 100% methanol, and stored at −20°C until being processed for WISH. Permeabilization incubation time in 10 µg/ml Proteinase K was optimized to maximize the signal in inner ear or posterior lateral line, with a 1-3 min step depending on the stage (3-5 dpf). Hybridization was performed at 70°C. Stained embryos were stored in 4% PFA until imaging using a ZEISS Axio Zoom V16 stereo microscope.

### Histology

After WISH staining, 5-day-old embryos were prepared for embedding in paraffin, transferred to paraffin, oriented, sectioned in ~5 µm transversal sections, and then counterstained using nuclear fast red (Sigma-Aldrich, cat. # N3020). Representative pictures were collected using an upright Zeiss Axio Imager D2 microscope (Carl Zeiss Inc.) with a Plan-Apochromat 20X/0.75 NA or a dry Plan-Neofluar 40X/0.75 NA objective lenses. All images were acquired using a AxioCam HRc full color CCD camera with a 1388 pixel x 1040 pixel imaging field. Zeiss ZEN blue pro 2011 software package was used for collection.

### Alkaline phosphatase staining

Alkaline phosphatase staining to visualize the posterior lateral line neuromasts was performed as previously described(*34*). Briefly, after over-night fixation in 4% PFA, the embryos were washed several times in PBS-Tween and then transferred to WISH staining buffer. The development of the signal was periodically checked under a stereomicroscope and it was stopped by washing with PBS-Tween and finally transferring the embryos in 4% PFA. Stained embryos were stored in 4% PFA until imaging using a ZEISS Axio Zoom V16 stereo microscope.

### Inner ear dissection and hair cell bundles staining

Inner ear dissections and phalloidin staining of hair cell bundles was performed as previously described(*70*). Briefly, after inner ear fixation in 4% PFA the different compartments of the inner ear (saccule, lagena and utricle) were dissected under a stereomicroscope and then stained with Alexa-labeled phalloidin (Alexa Flour 488 Phalloidin, 1:1,000 dilution, Thermo Fisher Scientific) for 30 min at room temperature to visualize hair cell bundles from *ak2^hg16/+^* and *ak2^hg16^* fish. Nuclei were stained with DAPI (Thermo Fisher Scientific, cat. #D1306). After the staining, the tissues were washed in PBS-Tween 3 times for 10 min each and then mounted onto a slide with mounting medium. Fluorescent signal was analyzed using a Zeiss LSM 880 confocal system (see below). Quantification was performed selecting representative 2500 µm^2^ areas for every structure (6 different areas for utricles, 4 in the case of saccules and lagenas) and the number of phalloidin positive bundles in each area was counted. Then the average number was calculated, and the statistical analysis was performed (see below)(*70*, *71*).

### Confocal imaging

Confocal images were collected at room temperature using a Zeiss LSM 880 confocal system fitted with an Airyscan module, mounted on an inverted Zeiss Axio Observer Z1 microscope with a Plan-Apochromat 20X/0.75 NA objective lens (Carl Zeiss Inc., Thornwood, NY, USA). Embryos were anesthetized and embedded in 1% low-melting agarose and imaged. All images were acquired in LSM 880 mode using a 32-channel GaAsP-PMT detector. Excitation wavelength of 488nm, 561nm and 405nm lasers were used for green, red and DAPI signals. A range of z-slices at optimal intervals was used depending on the zebrafish orientation to capture all desired structures. Z-sections were collected at defined intervals, and maximum projections were performed on each Z-stack. All confocal images were of frame size 512 by 512 or 1024 by 1024 pixels, scan zoom of 0.6 and line averaged 2 times. Zeiss ZEN 2.3 (black) software package was used for collection and post processing of the images (Carl Zeiss).

### Hair cell ablation and regeneration assay

Hair cells were ablated at 3, 4, or 5 dpf by treatment with 10 µM copper(II) sulfate (Sigma-Aldrich, cat #451657) in 1x Holt’s buffer for 1 hour(*66*). After hair cell ablation, the embryos were kept in fresh Holt’s buffer for two days to allow recovery and hair cell regeneration. Functional hair cells of neuromasts were stained using 2 µM solution of Yo-PRO-1 iodide (Thermo Fisher Scientific, cat. #Y3603) in 1x Holt’s buffer for 1 hour. Stained embryos were washed three times with 1x Holt’s buffer and then anesthetized with 0.02% w/v MS-222 (Sigma-Aldrich, cat. #A5040) before the analysis. The embryos were then transferred to a 96-well plate (1 embryo/well) and orientated to a lateral position for counting positive hair cells in the lateral line neuromasts. Positive mature hair cells in P1-P4 lateral line primary neuromasts or in LII.1-LII.2 secondary neuromasts were counted using a Zeiss Axio Observer A1 Inverted Phase Contrast Microscope and then the average number of the regenerated hair cells was calculated(*72*, *73*). To confirm the correct ablation of hair cells, a small number of copper-treated embryos were stained using Yo-PRO-1 iodide immediately after the copper treatment.

### TUNEL assay

TUNEL assay to label apoptotic and dead cells was performed using the ApopTag® Red In Situ Apoptosis Detection Kit (Millipore, cat. #S7165) following the manufacturer’s instructions with minor modifications. Permeabilization incubation time in 10 µg/ml Proteinase K was optimized to maximize the signal in inner ear or posterior lateral line as was done in WISH experiments (1-3 min max depending on the stage).

### Chemical treatments

Zebrafish embryos were exposed to different doses of l-Glutathione reduced (GSH) (Sigma-Aldrich, cat. # G4251) from 24 hpf until 5 dpf in E3 embryo medium containing 0.003% PTU. New embryo medium with fresh compound was administered daily until 5 dpf. AG1478 (used 2-3 µM, Calbiochem, cat. # 658552) and 6-BIO (0.2 µM, Sigma-Aldrich, cat. #B1686) compounds or the same amount of DMSO (Sigma-Aldrich, cat. #D2650) used as control were added to embryo media from 2 to 5 dpf.

### Statistical analysis

The statistical analysis was conducted using two-tailed Student’s t-test using Excel (Microsoft) or a one-way ANOVA analysis followed by Tukey’s post hoc test (p<0.001) to compare groups of fish treated differently using PRIZM (GraphPad, La Jolla, CA). A difference was considered as significant when the p-value was less than 0.05 for Student’s t-test and less than 0.001 for the Tukey’s post hoc test. Asterisks refer to significant p-values. Bar graphs showed the mean and the standard errors or the standard error of the mean depending on the test used. All experiments shown were replicated at least two times.

## Supporting information

primers used in study

ak2 mutant primII migration

wild-type primII migration

## Acknowledgements

This research is funded by the Intramural Research Program of the National Human Genome Research Institute; National Institutes of Health (S.M.B.: 1ZIAHG000183). We would like to thank the staff of Charles River for animal care, Oreet Zimand for the help with drug treatment and WISH experiments, Dr. Erica Bresciani and Dr. Wuhong Pei for helpful discussions and suggestions, and all other members of the Burgess Lab.

---

**Figure 1-figure supplement 1.**
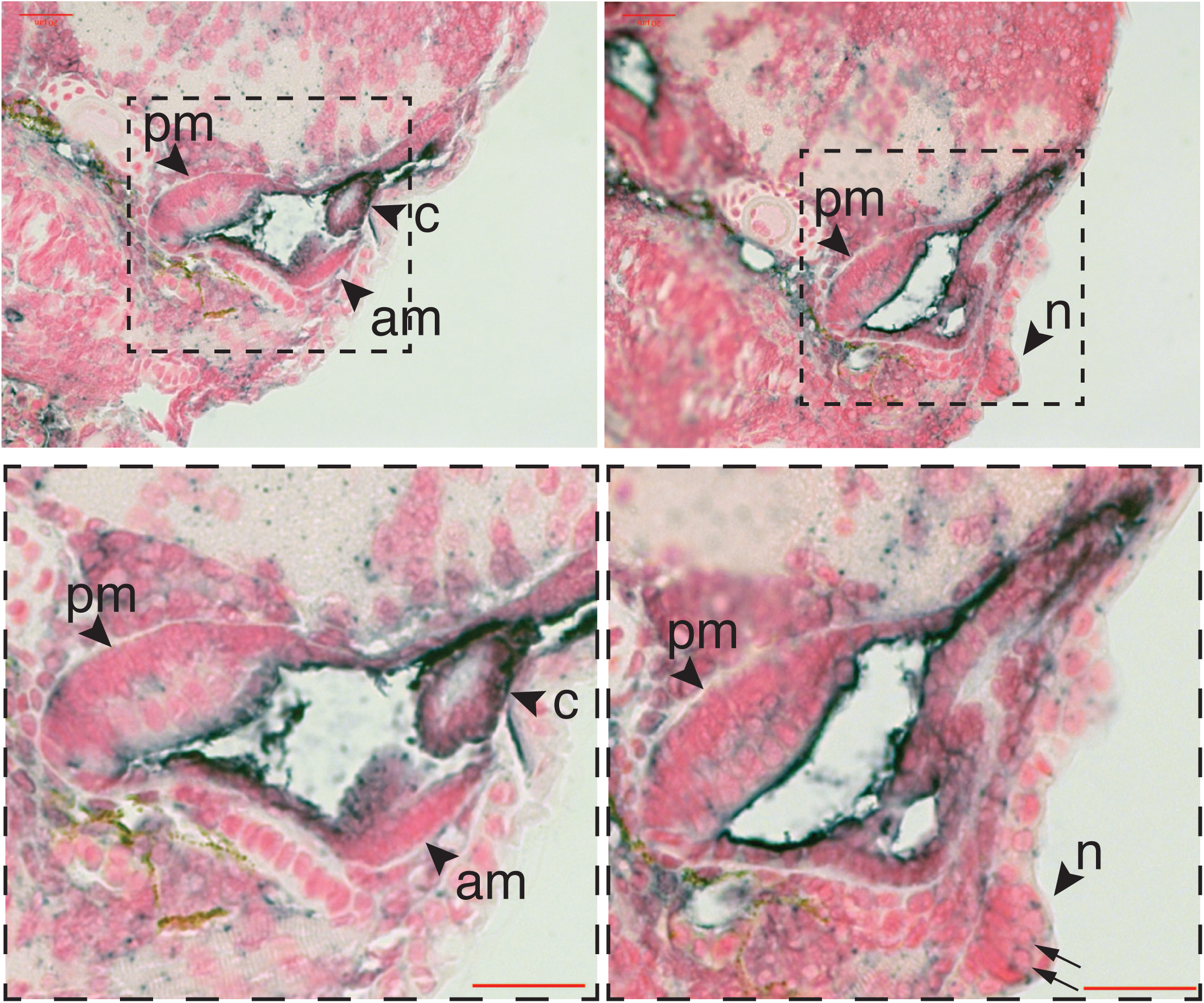
*ak2* expression in the otic vesicle and neuromasts in 5 dpf zebrafish larvae. Paraffin transverse sections through otic vesicle of 5 dpf wild-type larvae stained with an *ak2* WISH probe (purple). Nuclei are labeled with nuclear fast red. Magnification: 40x. Insets show higher magnifications of the same dashed areas. Black arrows in insets point to hair cells inside the neuromast structure. Scale bars: 20 µm.

**Figure 1-figure supplement 2.**
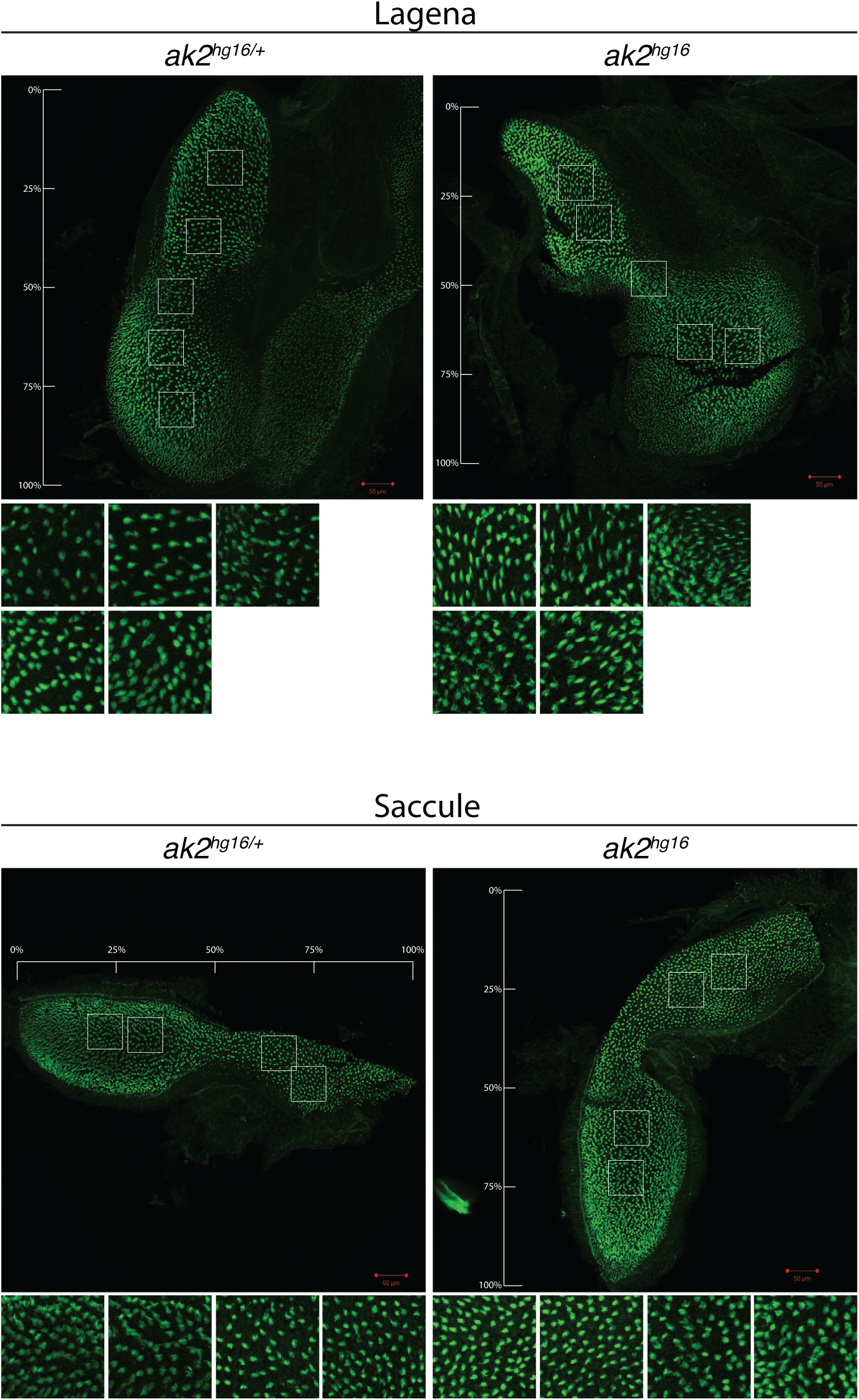
Comparison between hair cell bundles density in lagenas and saccules of *ak2^hg16/+^* and *ak2^hg16^* adult fish. The hair cell bundles were stained with phalloidin (green), nuclei with DAPI (blue). In each structure, hair cell bundle counts were sampled at 2500 µm^2^ areas indicated by white boxes and insets below each figure. Scale bars: 50 µm.

**Figure 2-figure supplement 1.**
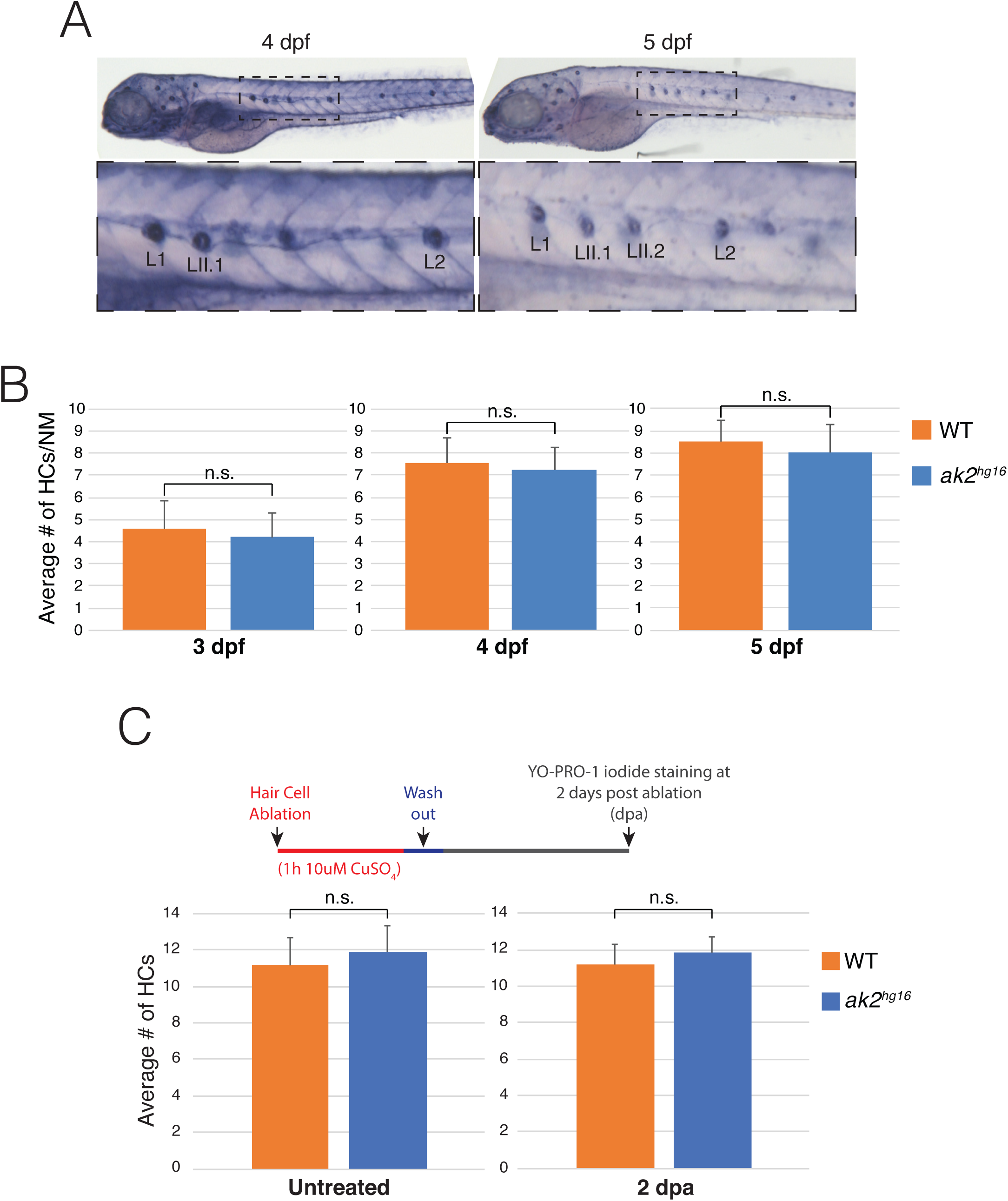
*ak2^hg16^* mutants show no detectable phenotypes in the lateral line neuromasts. (A) Alkaline phosphatase staining was performed on 4 and 5 dpf embryos obtained from an incross of *ak2^hg16^*^/+^. No differences were observed compared to control embryos (see Figure 2A for a refence of AP staining on WT embryos). Lateral views with anterior to the left. Dashed lines indicate insets of trunk regions. (B) Hair cells of lateral line neuromasts were fluorescently labeled with Yo-PRO-1 iodide at different stages to compare the average number of HCs per neuromast in *ak2^hg16^* and control embryos. (C) Analysis of hair cell regeneration after ablation using CuSO_4_ treatment in *ak2^hg16^* embryos and their siblings. n.s. denotes not significant values according to Student’s t-test.

**Figure 2-figure supplement 2.**
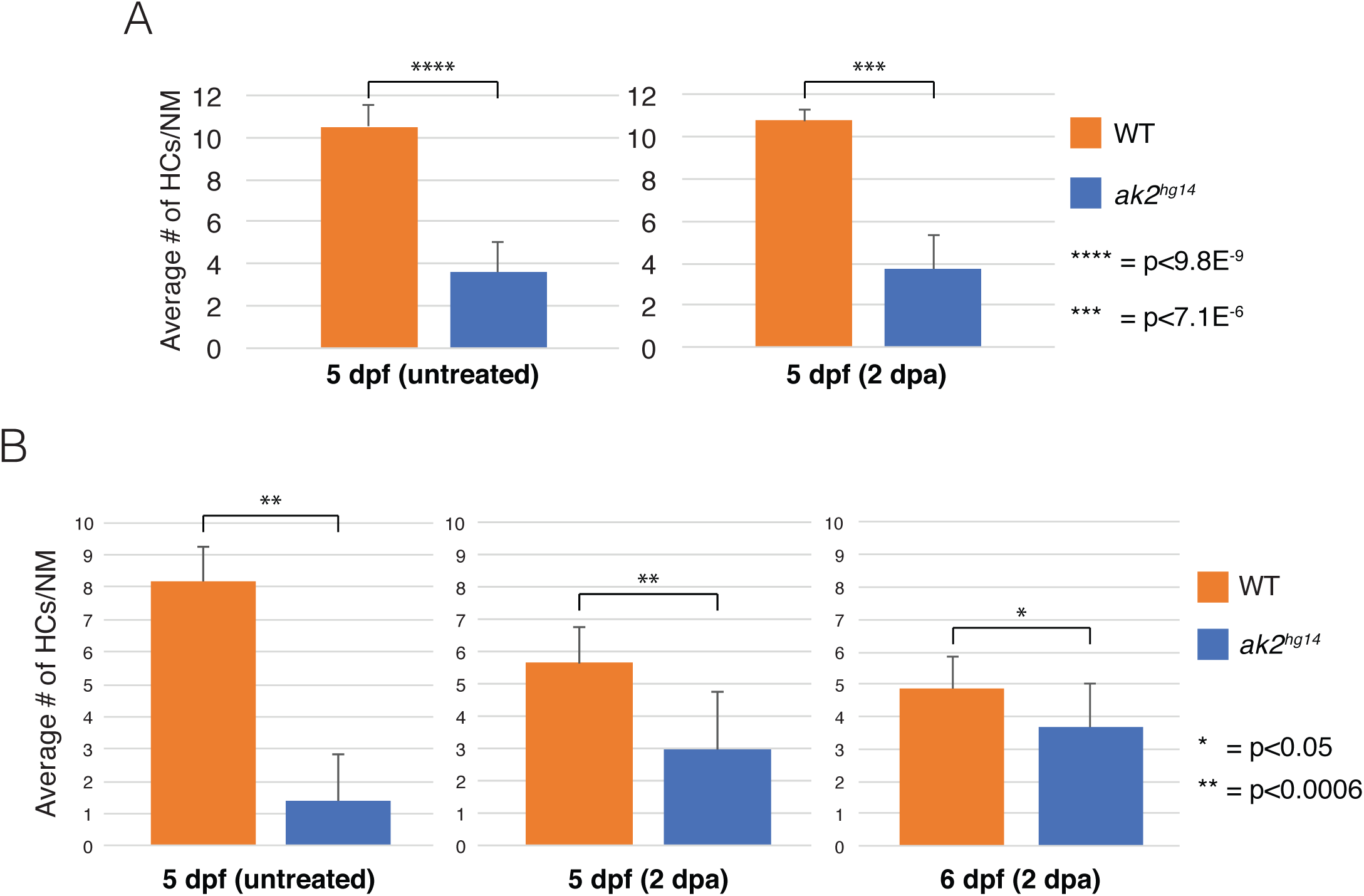
Hair cell regeneration after CuSO_4_ treatment in *ak2^hg14^* transgenic mutants and their siblings. (A) Analysis of the average number of GFP positive mature HCs per neuromast in Tg(*pou4f3*:GAP-GFP) *ak2^hg14^* embryos and their siblings after 2 days post ablation (dpa) with CuSO_4_. (B) Analysis of the average number of dTOM^+^ immature HCs per neuromast in Tg(*tnks1bp1*:EGFP; *atoh1a*:dTOM) *ak2^hg14^* embryos and their siblings after 2 days post ablation (dpa) with CuSO_4_. The error bars indicate standard errors. The asterisks denote significant values according to Student’s t-test. p indicates the p-values.

**Figure 3-figure supplement 1.**
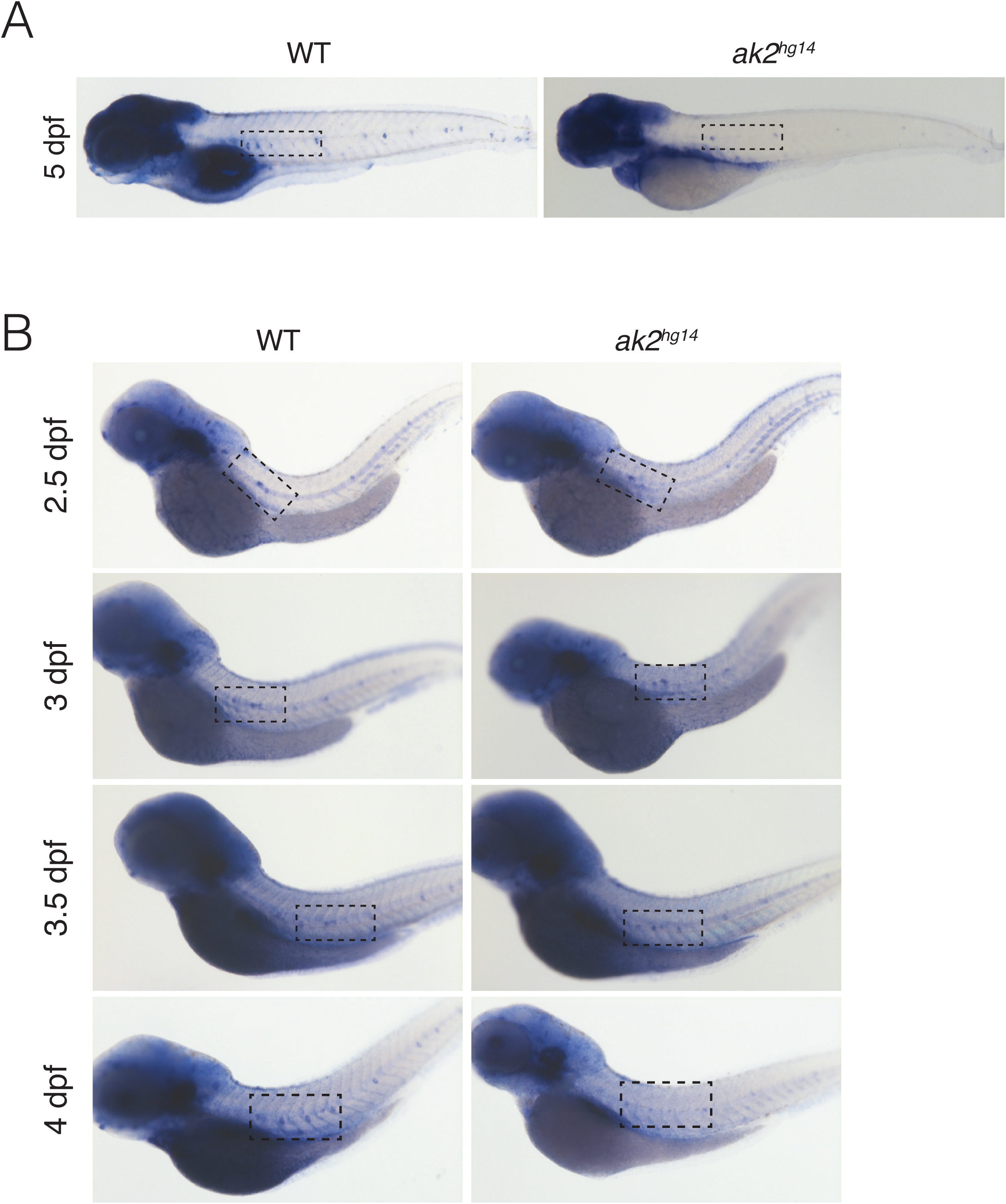
Characterization of *atoh1a* and *eya1* expression in the lateral line of *ak2^hg14^* mutants. (A) Expression of the *atoh1a* marker in primary and secondary neuromasts on 5 dpf *ak2^hg14^* embryos and their control siblings. Dashed insets indicate the trunk regions shown in Figure 3A. (B) *eya1* gene expression analysis in *ak2^hg14^* and control embryos at different stages of development (2.5 to 4 dpf). Dashed insets indicate the trunk regions shown in Figure 3C.

**Figure 6-figure supplement 1.**
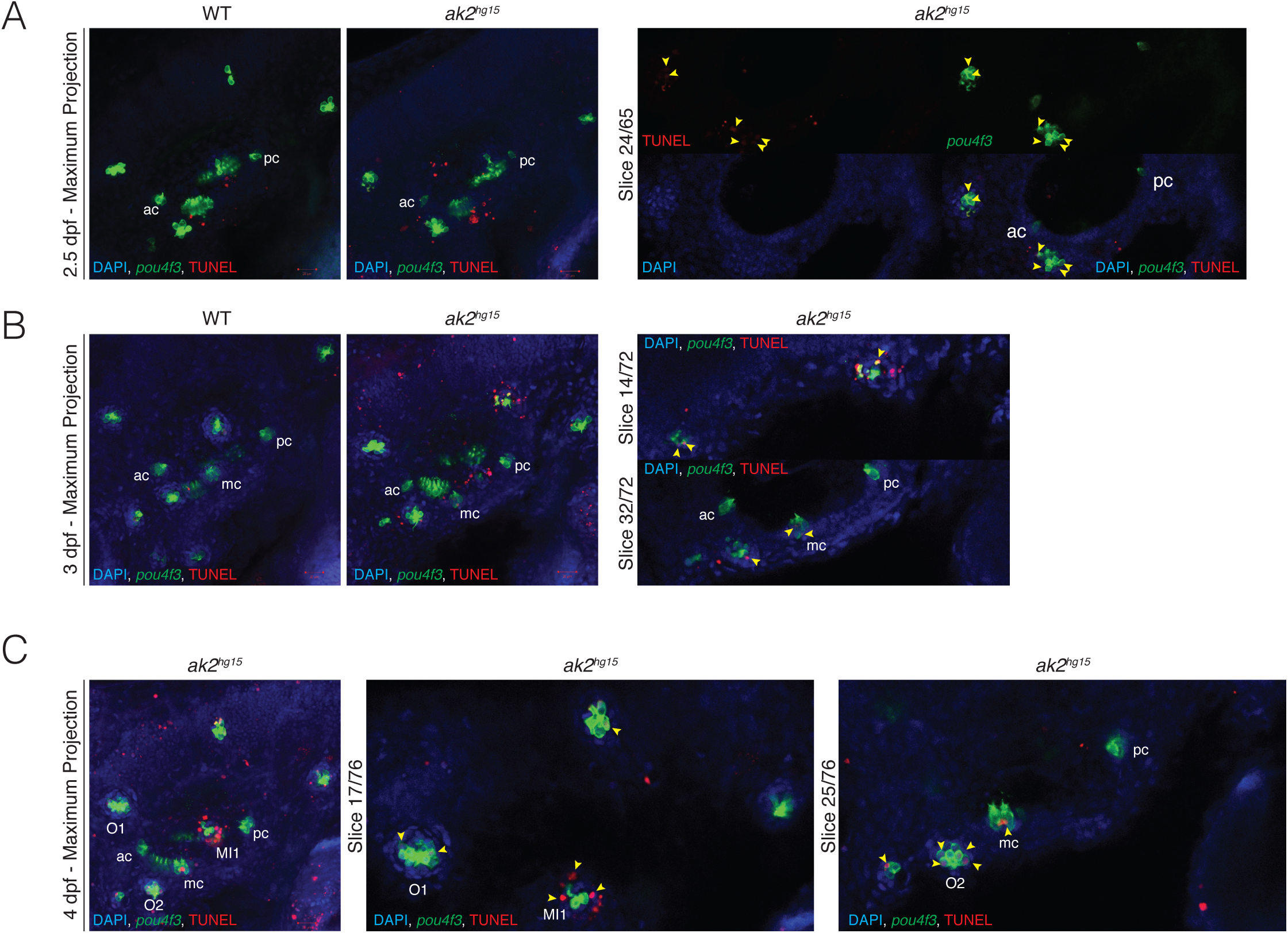
TUNEL staining in the otic vescicles of *ak2^hg15^* embryos. (A-B) Maximum projection (left panels) and representative single plane (right panels) confocal analyses at 2.5 and 3 dpf of TUNEL assays (red signal) performed on ak2*^hg15^* embryos and their control siblings in the Tg(*pou4f3*:GAP-GFP) background (indicated as *pou4f3*). Yellow arrowheads indicate examples of TUNEL positive cells. Tg(*pou4f3*:GAP-GFP) line labels the mature HCs (green), nuclei are labelled with DAPI (blue). (C) Maximum projection (left panels) and representative single plane (right panels) confocal analysis at 4 dpf of a TUNEL assay (red signal) performed on ak2*^hg15^* embryos in the Tg(*pou4f3*:GAP-GFP) background (indicated as *pou4f3*). Yellow arrowheads indicate examples of TUNEL positive cells. Tg(*pou4f3*:GAP-GFP) line labels the mature HCs (green), nuclei are labelled with DAPI (blue). ac: anterior crista; mc: medial crista; pc: posterior crista. Anterior lateral line nomenclature: otic (O1, O2) and middle (MI1) neuromasts. Scale bars: 20 µM.

**Figure 7-figure supplement 1.**
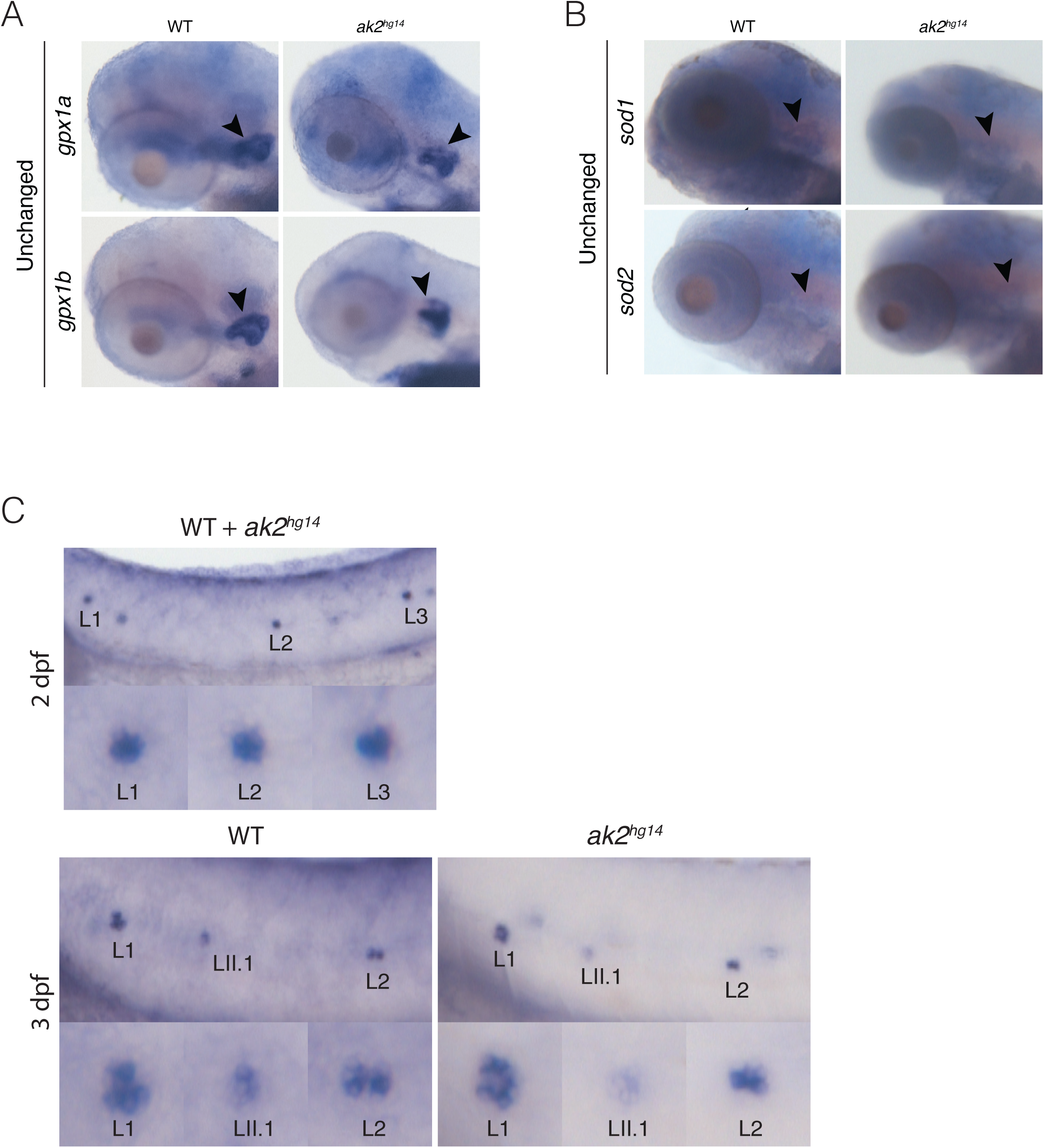
WISH analysis of oxidative stress markers in *ak2^hg14^* embryos. In situ hybridization analysis of *gpx1a* and *gpx1b* markers at 4 dpf (A), *sod1* and *sod2* markers at 5 dpf (B). The analysis showed no differences in the expression of the markers. (C) WISH analysis of *gstp1* gene expression at 2 and 3 dpf. L1-L3 define primary neuromasts; LII.1 designates the lateral line secondary neuromasts. Lateral views with anterior to the left. Black arrowheads indicate the otic vescicle. *gpx*: glutathione peroxidase; *sods*: superoxide dismutase; *gstp1*: glutathione S-transferase pi 1.

**Figure 8-figure supplement 1.**
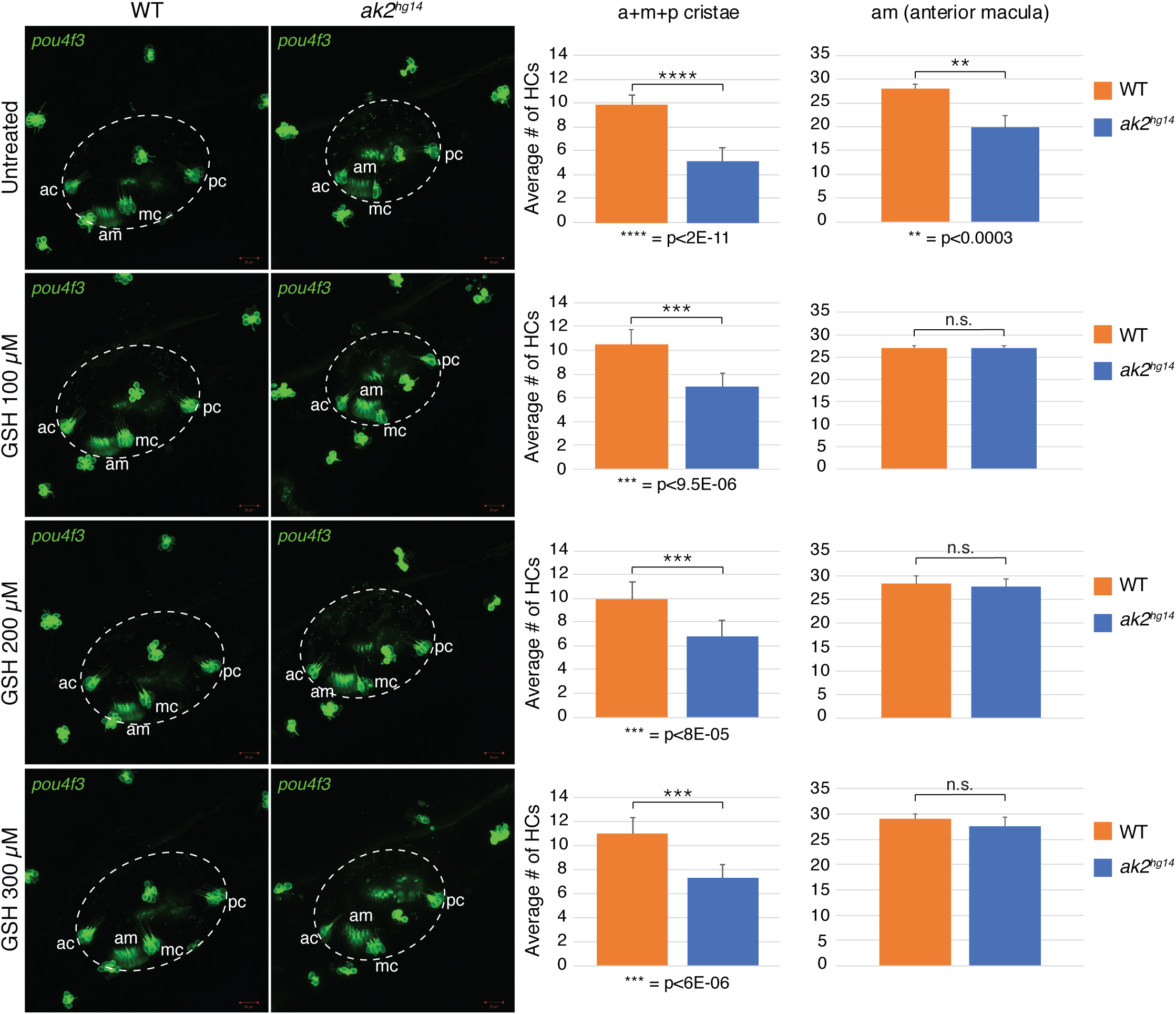
Confocal analysis of the inner ear region of *ak2^hg14^* mutants and their control siblings treated with different concentrations of glutathione until 3 dpf. Confocal maximum projections of the inner ear region (dashed circles) of 3 dpf Tg(*pou4f3*:GAP-GFP) *ak2^hg14^* mutants and control siblings, treated with different doses of GSH (100-300 µM). Graphs on the right of each treatment show the comparison of the average number of GFP^+^ hair cells in the cristae (a+m+p) and the anterior macula (am). The error bars indicate standard errors; asterisks denote statistically significant values. p indicates the p-value according to a Student’s t-test. A p<0.05 indicates a statistically significant effect. N.s.: not significant according to Student’s t-test. Magnification: 30x, scale bars: 20µm. Lateral views with anterior to the left. *pou4f3* denotes the Tg(*pou4f3*:GAP-GFP) line labelling the mature hair cells (green). ac: anterior crista; mc: medial crista; pc: posterior crista; am: anterior macula.

**Figure 8-figure supplement 2.**
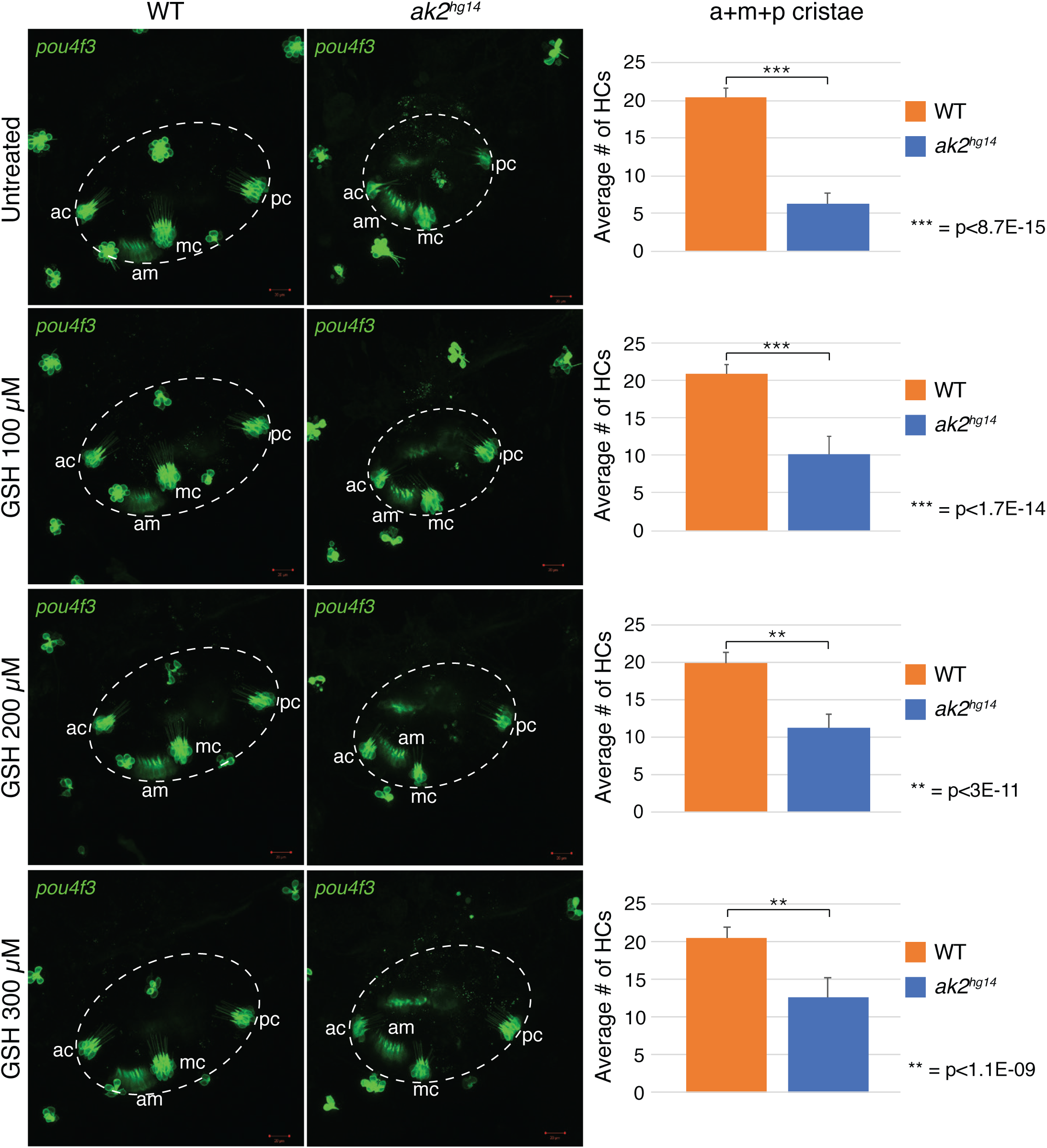
Confocal analysis of the inner ear region of *ak2^hg14^* mutants and their control siblings treated with different concentrations of glutathione until 4 dpf. Confocal maximum projection of the inner ear region (dashed circles) of 4 dpf Tg(*pou4f3*:GAP-GFP) *ak2^hg14^* mutants and control siblings, treated with different concentrations of GSH (100-300 µM). Magnification: 30x; scale bars: 20µm. Graphs on the right of each treatment show the comparison of the average number of GFP^+^ hair cells in the cristae (a+m+p) at 4 dpf between *ak2^hg14^* null and control embryos. Lateral views with anterior to the left. *pou4f3* denotes the Tg(*pou4f3*:GAP-GFP) line labelling the mature hair cells (green). ac: anterior crista; mc: medial crista; pc: posterior crista; am: anterior macula. The error bars indicate standard errors; asterisks denote statistically significant values; p indicates the p-value according to a Student’s t-test. A p<0.05 indicates a statistically significant effect.

Video 1.

Time-lapse recording of the migration of primII in an *ak2^hg15^* Tg(−8.0*cldnb*:LY-EGFP) mutant embryo from ~3.5 to ~4 dpf. Each frame was taken every ~10 min.

Video 2.

Time-lapse recording of the migration of primII in a control Tg(−8.0*cldnb*:LY-EGFP) embryo from ~3.5 to ~4 dpf. Each frame was taken every ~10 min.

